# Comparing Raman and NanoSIMS for heavy water labeling of single cells

**DOI:** 10.1101/2024.07.05.602271

**Authors:** George A. Schaible, John B. Cliff, Jennifer A. Crandall, Jeremy J. Bougoure, Michael N. Mathuri, Alex L. Sessions, Joseph Atwood, Roland Hatzenpichler

**Affiliations:** Department of Chemistry and Biochemistry, Montana State University, Bozeman, MT 59717; Center for Biofilm Engineering, Montana State University, Bozeman, MT 59717; Environmental Molecular Sciences Laboratory, Pacific Northwest National Laboratory, Richland, WA 99354; Department of Microbiology and Cell Biology, Montana State University, Bozeman, MT 59717; Division of Geological and Planetary Sciences, California Institute of Technology, Pasadena, CA 91125; Department of Agricultural Economics and Economics, Montana State University, Bozeman, MT 59717; Thermal Biology Institute, Montana State University, Bozeman, MT 59717

**Keywords:** Cellular heterogeneity, D_2_O, heavy water, cell activity, single cell microbiology, technique comparison, Raman, NanoSIMS

## Abstract

Stable isotope probing (SIP) experiments in conjunction with Raman microspectroscopy (Raman) or nano-scale secondary ion mass spectrometry (NanoSIMS) are frequently used to explore single cell metabolic activity in pure cultures as well as complex microbiomes. Despite the increasing popularity of these techniques, the comparability of isotope incorporation measurements using both Raman and NanoSIMS directly on the same cell remains largely unexplored. This knowledge gap creates uncertainty about the consistency of single-cell SIP data obtained independently from each method. Here, we conducted a comparative analysis of 543 *Escherichia coli* cells grown in M9 minimal medium in the absence or presence of heavy water (^2^H_2_O) using correlative Raman and NanoSIMS measurements to quantify the results between the two approaches. We demonstrate that Raman and NanoSIMS yield highly comparable measurements of ^2^H incorporation, with varying degrees of similarity based on the mass ratios analyzed using NanoSIMS. The ^12^C^2^H/^12^C^1^H and ^12^C_2_^2^H/^12^C_2_^1^H mass ratios provide targeted measurements of C-H bonds but may suffer from biases and background interference, while the ^2^H/^1^H ratio captures all hydrogen with lower detection limits, making it suitable for applications requiring comprehensive ^2^H quantification. Importantly, despite its higher mass resolution requirements, the use of C_2_^2^H/C_2_^1^H may be a viable alternative to using C^2^H/C^1^H due to lower background and higher overall count rates. Furthermore, using an empirical approach to determining Raman wavenumber ranges via the 2^nd^ derivative improved the data equivalency of ^2^H quantification between Raman and NanoSIMS, highlighting its potential for enhancing cross-technique comparability. These findings provide a robust framework for leveraging both techniques, enabling informed experimental design and data interpretation. By enhancing cross-technique comparability, this work advances SIP methodologies for investigating microbial metabolism and interactions in diverse systems.

**Importance:** Accurate and reliable measurements of cellular properties are fundamental to understanding the function and activity of microbes. This study addresses to what extent Raman microspectroscopy and nano-scale secondary ion mass spectrometry (NanoSIMS) measurements of single cell anabolic activity can be compared. Here, we study the relationship of the incorporation of a stable isotope (^2^H through incorporation of ^2^H_2_O) as determined by the two techniques and calculate a correlation coefficient to support the use of either technique when analyzing cells incubated with ^2^H_2_O. The ability to discern between the comparative strengths and limitations of these techniques is invaluable in refining experimental protocols, enhancing data comparability between studies, data interpretation, and ultimately advancing the quality and reliability of outcomes in microbiome research.

## Introduction

Microbial activities and cellular interactions drive human health, global biogeochemical cycles, and ecosystem function. Stable isotope probing (SIP) is a particularly powerful approach to studying microbial function and is widely used in microbiology (1-5). By measuring the incorporation of heavy isotopes (*e*.*g*., ^2^H, ^13^C, ^15^N, ^18^O) into biomass, it is possible to assess the activity and physiology of individual cells at a microbially-meaningful spatial resolution, enabling a broader understanding of community dynamics (4). While SIP can be applied to all assimilatory pathways, heavy water labeling holds appeal for microbiologists due to its universal applicability, negating the need for prior knowledge of cell physiology (2). When combined with the addition of unlabeled substrates, heavy water labeling enables researchers to pinpoint which cells within complex microbiomes are activate in presence of a substrate (1). Active microbes grown or incubated in the presence of heavy water will use deuterium (^2^H) in lieu of hydrogen (^1^H) during biosynthesis, resulting in active microbial community members becoming labeled with ^2^H (6, 7). Because ^2^H naturally occurs at very low levels (∼0.016%), its environmental background is usually negligible for biological labeling studies (8, 9).

The assimilation of isotopically labeled substrates, including heavy water, into individual cells can be measured by either Raman microspectroscopy (Raman) or nano-scale secondary ion mass spectrometry (NanoSIMS). Micro-autoradiography is a cost-effective alternative but is limited to radioactive isotopes (10, 11). All three techniques can be combined with fluorescence *in situ* hybridization (FISH), which allows a direct link between cellular identity and function in complex microbiomes (3, 12-14).

Raman is a non-destructive spectroscopic technique based on the inelastic scattering of light (3, 15). It can be applied directly to environmental samples with minimal preparation (16), allowing for real-time monitoring of metabolic changes of single cells (17). By exciting molecular bond vibrations, Raman elucidates the spatial distribution of specific biomolecules, such as lipids, proteins, and nucleic acids within cells and/or communities on a sub-micrometer scale (3, 16). Incorporation of stable isotopes into biomolecules changes the vibrational energy of excited molecules through the increased molecular mass, inducing a red shift towards smaller wavenumbers (16, 18). While Raman requires little or no sample preparation, the technique suffers from low sensitivity (∼10% ^13^C, ∼10% ^15^N, and ∼0.2% ^2^H cellular replacement (16, 18, 19)) and relatively low signal-to-noise ratio as compared to mass spectrometry. However, carbon-deuterium (^12^C^2^H) bonds are reliably detected in Raman spectra by a characteristic peak shift of the abundant ^12^C^1^H peak into the mostly Raman-silent region (1,800-2,800 cm^-1^) of the cellular spectrum making Raman analysis well suited for deuterium SIP analysis in single cells (2).

NanoSIMS uses a focused ion beam to produce secondary ions from a sample surface that are subsequently analyzed via a double focusing mass spectrometer to attain spatially resolved elemental and isotopic information (20, 21). Although NanoSIMS is capable of detecting isotopes at a much greater sensitivity and spatial resolution (≥50 nm) than Raman, it is destructive to the sample and provides limited molecular information (21). Furthermore, unlike Raman, the molecular pools of H that the NanoSIMS interrogates is unclear. NanoSIMS can suffer from recombination of atoms during ion beam sputtering, which may result in misinterpretation of results. However, in the case of ^13^C- and ^15^N-labeling of biological materials, recombination is not thought to affect the general interpretation of NanoSIMS results (22). At the whole-cell level, this assumption likely holds for ^2^H as well. However, as this study highlights, heterogeneous sampling of cellular components and the choice of ion pair complicate detailed data interpretation with our current understanding.

Both Raman and NanoSIMS are powerful tools for studying the structure, function, and interactions of microbes on a single-cell or community level (3, 21). Studies have successfully applied both techniques in tandem, including analysis of deposition of organic material in diatom fossils (23), SIP experiments of microbial cells recovered from soils and sediments (24, 25), and *in vitro* SIP of microbial cells for sorting (2). However, to the best of our knowledge, only two studies ever applied Raman and NanoSIMS on the same cells (25, 26), and a comprehensive and direct comparison of these techniques was previously unavailable, making it difficult to compare and interpret results generated by these techniques.

To address this shortcoming, we employed SIP on *Escherichia coli* using deuterium oxide (^2^H_2_O) and compared results of Raman and NanoSIMS based measurements of 543 individual cells. We demonstrate strong correlation of these measurements, enabling the corroboration of results from both techniques. This data set also enabled an investigation into the pre-processing of Raman data providing insights on how to best optimize the methodology when using either technique for ^2^H incorporation measurements. In addition, we investigated a statistical approach for calculating wavenumber ranges for ^12^C^2^H measurements using Raman.

Considering that both Raman and NanoSIMS possess unique strengths and limitations, the drawbacks of one technique can be compensated for by the other, thereby granting a more comprehensive insight into biological samples at a micrometer scale (25). Ultimately, this comparative investigation contributes to refining our understanding of Raman and NanoSIMS data, lays the foundation for future integrated approaches of correlative SIP-Raman-NanoSIMS analyses, and illuminates the strengths and weaknesses of each technique to allow future investigators more robust information when evaluating the applicability of both techniques for a given experimental objective.

## Results and Discussion

### Stable isotope probing

We employed our recently developed correlative workflow (25) for the study of single *E. coli* K12 cells that had been grown in unamended media (0% added ^2^H_2_O) or media amended with either 15%, 30%, or 50% ^2^H_2_O. Lower ^2^H_2_O amounts (between 0% and 15%) were not tested due to the limited sensitivity of Raman spectroscopy for detecting ^2^H_2_O uptake at low enrichment levels (25). In such cases, NanoSIMS is the preferred method, because it offers much higher sensitivity and thus enables detection of low levels of anabolic activity (27). Cells were collected during mid-logarithmic growth (Fig. 1 and S1), fixed with paraformaldehyde (PFA) to preserve cells and inhibit change in ^2^H content, immobilized on a stainless-steel coupon, stained with 4’,6-diamidino-2-phenylindole (DAPI, necessary for visualization during next step), and regions of interest (ROIs) mapped using laser dissection microscopy (Fig. S2). Raman spectra were collected for each individual cell within the defined ROIs before NanoSIMS measurements were performed.

**Fig. 1.**
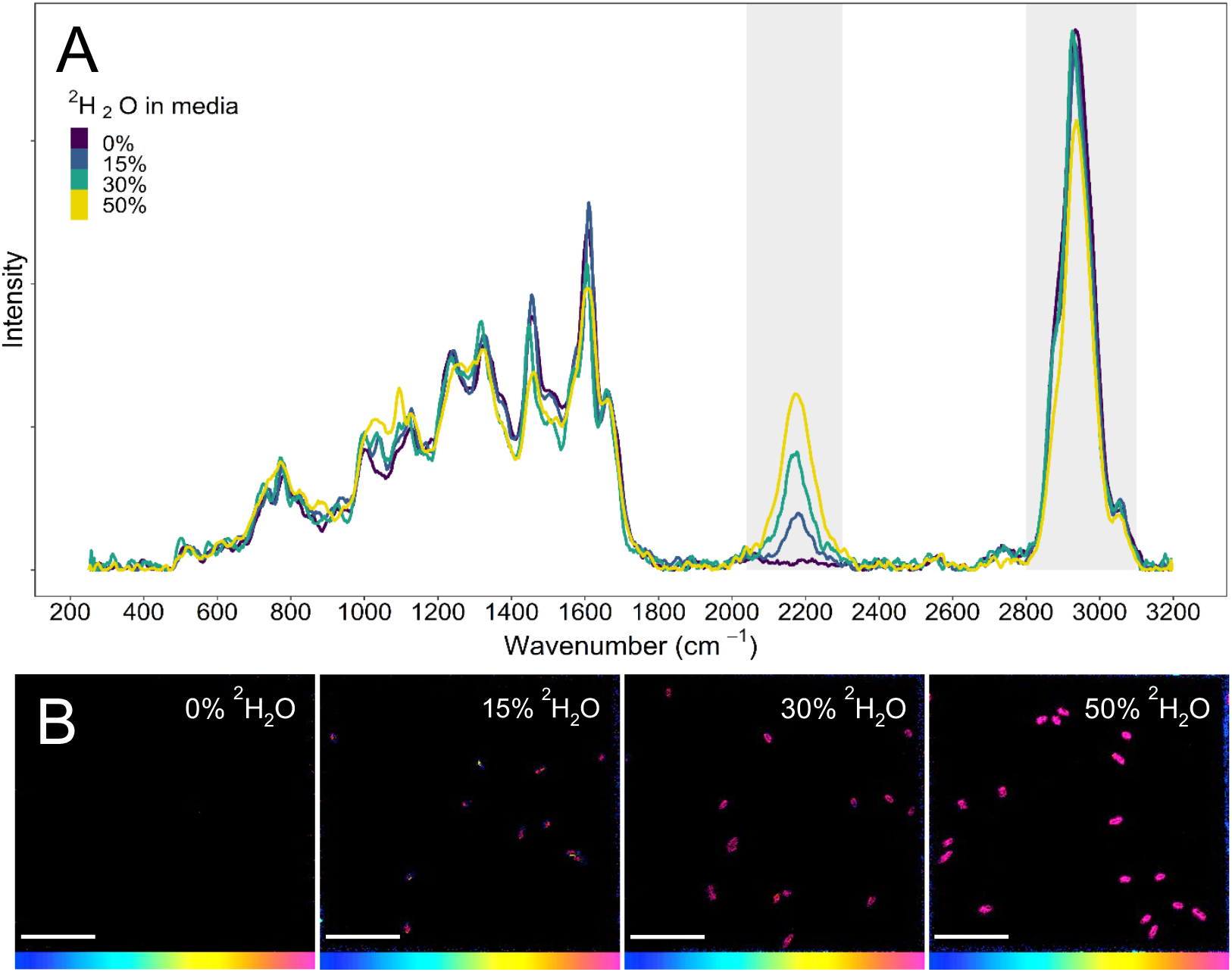
Representative Raman spectra and NanoSIMS images illustrating the uptake of ^2^H-label by *Escherichia coli* K12 cells after incubation in media with varying percentages of ^2^H_2_O. **A** The characteristic ^12^C^2^H and ^12^C^1^H regions are shaded in gray. Each Raman spectrum was obtained from a single cell. **B** ^12^C_2_^2^H/^12^C_2_^1^H mass ratio NanoSIMS images. All isotope fraction images are on the same scale (0-30 atom %). Scales bars are 10 µm.

### Validation of natural abundance using IRMS

Discrepancies between techniques for ^2^H measurements for 0% ^2^H_2_O incubations prompted additional analysis to confirm the natural abundance of ^2^H in these samples. To investigate this, we used isotope ratio mass spectrometry (IRMS) on lipids extracted from the 0% ^2^H_2_O incubation samples. The lipids were selected for IRMS analysis because they are reflective of the isotopic signature of cellular material without being influenced by extracellular artifacts. Extracted lipids were esterified to form fatty acid methyl-esters (FAMEs) for analysis, enabling better ionization and separation during IRMS (28). The analysis showed that lipids from the 0% ^2^H_2_O incubations were at natural ^2^H abundance as compared to the VSMOW international standard (Table S1). This finding confirmed that the elevated ^2^H levels observed in Raman and NanoSIMS 0% ^2^H_2_O data were due to conflated values rather than true biological enrichment. Furthermore, previous studies comparing bulk elemental analyzer IRMS measurements to SIMS have shown that IRMS values are generally higher than those derived from SIMS (29), further supporting that the 0% ^2^H_2_O incubations did not contain any incorporated ^2^H.

### Pre-processing of Raman spectra

Currently, there is no standard method for the pre-processing of Raman spectra (16), despite compelling evidence demonstrating the influence of processing procedures on data outcomes (30). Optimizing the workflow by testing various pre-processing procedures on Raman spectra before selecting a method is recommended to minimize information loss in the spectra (31, 32). To address the lack of standardization in pre-processing methods, we systematically tested various methods on our Raman spectra to optimize our spectra processing methodology. Our findings showed that pre-processing techniques had little effect on the quantification of ^2^H from *E. coli* spectra.

Raman spectra spanning 250-3,200 cm^-1^ were acquired from individual cells within the ROIs. To test if peaks in the fingerprint region of spectra (ca. 700-1,800 cm^-1^) induce a leveraging effect during smoothing and baselining of the spectra, spectra were analyzed as a whole (250-3,200 cm^-1^) or were truncated to 1,800-3,200 cm^-1^, as was previously suggested (31, 33). All spectra were smoothed using a Savitzky-Golay smoothing algorithm, a widely used technique for spectroscopy data (16, 32). This algorithm applies a moving window with a local polynomial fitting procedure to reduce noise while preserving peak shapes (34). It has been shown that excessive smoothing of spectra can result in the loss of genuine Raman peaks, thereby producing inaccurate spectra (30). To avoid this, we used a 2^nd^ degree polynomial with a window size of 11, as these settings preserved important peaks across all spectra. Next, several commonly applied baselining methods were tested to identify if background correction influences the final calculation of ^2^H content of the *E. coli* cells.

The first baselining method applied was asymmetric least squares (ALS), an iterative method that uses asymmetric weighting to adjust the extent in which data influence baselining (35). The second method used was a polynomial fit, where a polynomial of n^th^ degree (we used a 2^nd^ degree polynomial) can be fit to both the spectra and the baseline subtracted to remove background (36). The third method tested was a frequency differentiated non-linear digital filter, commonly referred to as “rolling ball”, which is equivalent to rolling a ball with a given ball diameter (*i*.*e*., window size) below the spectra and removing the background below the points the ball touches (37). The fourth and final method tested was statistics-sensitive non-linear iterative peak-clipping (SNIP), a process that compresses the data using a log square root operator followed by iterative evaluations using a clipping window, after which the data is transformed back using the inverse log square root (38). In contrast to the other baselining methods, SNIP does not lead to distortions of the baseline at the edges of the spectral interval (31).

Prior to determining the efficacy of the baselining methods, each spectrum was normalized using sum normalization to maintain relative proportions of peaks for comparisons. Next, the ^2^H incorporation into cells was quantified by calculating the ratio between the area under the curve (AUC) of the ^12^C^2^H (2,040-2,300 cm^-1^) and the ^12^C^1^H (2,800-3,100 cm^-1^) region of each spectrum (Fig. 1A). The AUC was determined by drawing a linear baseline between the interval of wavenumbers and numerically integrating the area above the line and below the curve using the trapezoidal rule (Fig. S3). The ^2^H content was then used to compare the baselining methods for both truncated spectra (1,800-3,200 cm^-1^) and whole spectra (250-3,200 cm^-1^), and the r-squared value used to determine the fit of each model (Fig. S4).

There was no statistically significant difference between the truncated and whole spectra or between the baselining methods, except for spectra baselined using the rolling ball method (Fig. S5, Table S2). This indicates that the baselining method had little effect on ^2^H calculation from Raman spectra of *E. coli*. This could be due to the low background observed in a pure culture as compared to an environmental sample and the low natural fluorescence of *E. coli*, resulting in minimal background subtractions from the data. We elected to use the SNIP baselining method for our analysis because this method has minimal distortion on data at both ends of the spectrum and has previously been used in single cell SIP-Raman studies (2, 24, 39).

### Analysis of NanoSIMS ion pairs

The next step in our workflow was to estimate the ^2^H content of each cell that had previously been measured by Raman using NanoSIMS (Fig. 1B). Historically, analysis of ^2^H uptake in cells by NanoSIMS has been performed using either the mass ratio of ^2^H/^1^H (25, 40, 41), ^12^C^2^H/^12^C^1^H (2, 42), or ^16^O^2^H/^16^O^1^H (2, 43). Among these, H^-^ and C_2_H^-^ have relatively higher ion yields compared to CH^-^ (Fig. S6), and the ^2^H/^1^H ratio requires relatively lower theoretical mass resolving power (MRP) to acquire a clean signal compared to organic ion pairs, which are more susceptible to overlapping interferences (44, 45). Although the use of ^12^C^2^H/^12^C^1^H and ^12^C_2_^2^H/^12^C_2_^1^H in polymer films and charcoal has been explored using high MRP, these studies relied on a high primary beam current, relatively low spatial resolution, and/or nonstandard instrument configurations to achieve optimum results (44, 45). While MRP is useful for resolving overlapping peaks, abundance sensitivity provides a more relevant measure of interference between ion species. However, in multi-species peaks, both metrics are generally impractical to measure directly. Confounding this difficulty is the use of multiple detectors during NanoSIMS analysis. Indeed, because of a variety of physical factors, identical ion species usually have different measured MRPs when measured on different detectors. In this study, we compare the use of ^2^H/^1^H, C^2^H/C^1^H, and C_2_^2^H/C_2_^1^H to detect ^2^H_2_O incorporation at the single cell level. To achieve the required spatial resolution, we use a relatively small primary beam current and standard MRP. We show that mass separation based on the concept of MRP alone is not a strict requirement for obtaining meaningful NanoSIMS data. We discuss our findings in relation to subcellular chemical morphology and experimental objectives. Further discussion on NanoSIMS methodology, including theoretical mass resolving power, detector tuning, and sources of analytical variability, is provided in the Supplemental Text.

Two observations stand out from the ^2^H/^1^H measurements. First, the background associated with cells grown without ^2^H_2_O was significantly lower than measurements using the CH^-^ and C_2_H^-^ ion pairs despite the presence of significant H in the stainless steel wafers. The lower background can be attributed to the relative ease of resolving isobars, providing evidence that our assigned ROIs did not include excessive amounts of H signal from the instrument vacuum or the stainless steel substrate. Second, the slope of the regression line for the % ^2^H_2_O in the growth medium versus NanoSIMS ^2^H/^1^H data is significantly lower than the corresponding regressions for the other ion pairs (Fig. S7, Table S3). This difference is likely a convolution of different IMFs between ion pairs analyzed as well as a difference in H pool being accessed. Assuming a value for the natural abundance ^2^H/H of 0.000156 Vienna Standard Mean Ocean Water (VSMOW), the slope of the regression of ^2^H/^1^H versus the concentration of ^2^H_2_O in the growth media (in %) is consistent with an instrumental mass fractionation (IMF) of approximately -380 ‰ compared with the regressions for ^12^C^2^H/^12^C^1^H, and ^12^C_2_^2^H/^12^C_2_^1^H (Fig. S7). This fractionation is in the range of values reported for SIMS analyses in the past (46) and is offset by ∼ 170 ‰ from the average value for fatty acids analyzed by IRMS (Table S1). We do not know what contribution IMF had versus real variability of ^2^H in cellular constituents. We would expect δ^2^H to change with cellular constituent as a function of depth. Using a previously published value for sputter rate of *Bacillus* spores using a 16 keV Cs^+^ primary ion beam of ∼2.5 nm um^2^ pA^-1^ s^-1^, we estimate that the NanoSIMS data presented here were acquired after removing about 20 nm of cell material indicating cell wall and membrane material likely have been interrogated, however, these assertions should be verified using careful future experimentation (47). In a complex environment like a bacterial cell, biochemically different pools account for different δ^2^H isotope values are expected to be encountered as a function of analysis depth. Large H isotope fractionations in fungal, algal, and plant cellular components have been previously documented as has fractionation of microbial lipids based on metabolic pathway (48-50). Intriguingly, our analysis showed a statistically significant difference (p < 0.005) between ^2^H/^1^H estimates for each pair of mass ratios used to measure the ^2^H content of cells at each concentration of ^2^H_2_O in the media (Fig. S8). On average, the ^12^C^2^H^-^/^12^C^1^H^-^ ratio indicated the highest apparent relative ^2^H content in cells, the ^12^C_22_H^-^/^12^C_2_^1^H^-^ ratio indicated the second highest, and the ^2^H^-^/^1^H^-^ ratio showed the lowest. This discrepancy was likely not sample-specific as the ^12^C^2^H^-^/^12^C^1^H^-^ and ^12^C_22_H^-^ /^12^C_2_^1^H^-^ mass ratios were concurrently measured on the same cells. Rather, the differences were likely due to some combination of sampling different ^2^H populations associated with the individual cell, different IMF factors, and variable atomic mixing during the sputtering process (22). Removal of isobaric interferences, such as the subtraction of the^13^C^1^H^-^ peak from the m/z 14 peak, would lead to lower detection limits for the ^12^C^2^H^-^/^12^C^1^H^-^ analyses (2). It is likely that calibration using a suitable organic ^2^H/^1^H standard would improve accuracy for all ion pairs.

Because DAPI staining was necessary to visualize cells during etching of the sample surface with the laser dissection microscope used for cell mapping, we investigated its effect on ^2^H signal dilution. Addition of hydrogen-rich chemicals, such as the fixatives paraformaldehyde (PFA) and ethanol or DNA-staining chemicals such as DAPI and SYBR dyes, can result in a dilution of isotopes in cells (43). Comparing ^2^H-labeled cells with and without DAPI revealed an average 12.8% decrease in total ^2^H signal for DAPI treated cells (Fig. S9). This dilution is expected, as each DAPI molecule contains fourteen hydrogen atoms and binds to twelve base pairs along the minor groove in DNA (51). Given the approximately 4.6 Mb long chromosome of *E. coli* K12, DAPI staining could introduce an estimated 5.4 × 10^6^ additional ^1^H per chromosome copy. Although this number is likely inflated due to the specific binding of DAPI to adenine-thymine rich regions in the minor groove of DNA, which are not present in every full turn of the minor groove (51), this example serves to highlight the effect DAPI can have on single cell isotope measurements. A previous study demonstrated that both DAPI and SybrGreen DNA-stains have a statistically significant dilution effect on ^13^C and ^15^N isotopes (^2^H was not evaluated). The authors concluded that if environmental microbes with low metabolic activity are analyzed, pretreatment of cells (PFA-fixation, DNA-staining, etc.) should be limited as much as experimentally possible (43). In our work, we did not evaluate the ^2^H-dilution effect of DAPI on Raman measurements.

### Single-cell comparison of Raman and NanoSIMS

Due to the variations in apparent ^2^H content of cells between the different mass ratios, we made our NanoSIMS ion pair comparisons using all three mass ratios. A comparison of the Raman spectra (analyzed using the 2040-2300 cm^-1^ wavenumber range for ^2^H calculations) and NanoSIMS measurements (for ^2^H/^1^H, ^12^C^2^H/^12^C^1^H, and ^12^C_2_^2^H/^12^C_2_^1^H mass ratios) of 543 cells revealed a high degree of comparability between the two techniques (Fig. 2). The highest correlation of labeling percentages determined by Raman and NanoSIMS were found using the ^12^C_2_^2^H/^12^C_2_^1^H mass ratio (0.98 r-squared). The ^2^H/^1^H and ^12^C^2^H/^12^C^1^H mass ratios showed equal correlation (0.97 r-squared) to the Raman results although the NanoSIMS data for the ^2^H/^1^H had a much lower degree of spread for the 0% ^2^H_2_O incubations as compared to ^12^C^2^H/^12^C^1^H. Analysis of covariance (ANCOVA) was used to evaluate the relationship between the linear models, showing that each model for each specific mass was significantly different (F-value > 150, p-value < 2×10^−16^). Considering the slope of the models, each comparison had a slope >1, indicating that Raman estimated a higher amount of deuterium in the cells than NanoSIMS. This discrepancy was the greatest with the ^2^H/^1^H mass ratio for which the slope of the model was 1.5. Furthermore, the y-intercept was close to zero for ^2^H/^1^H and ^12^C_2_^2^H/^12^C_2_^1^H while the ^12^C^2^H/^12^C^1^H mass ratio had a y-intercept of -4.86, representative of the high estimation of ^2^H in the cells for the 0% ^2^H_2_O incubation. Comparison of ^2^H content in each cell based on ^2^H_2_O incubations as measured by NanoSIMS each or Raman revealed statistically significant differences between the techniques (Fig. 3). The ^12^C_2_^2^H/^12^C_2_^1^H mass ratio was not significantly different from the Raman data at the 0% and 15% ^2^H_2_O incubations.

**Fig. 2.**
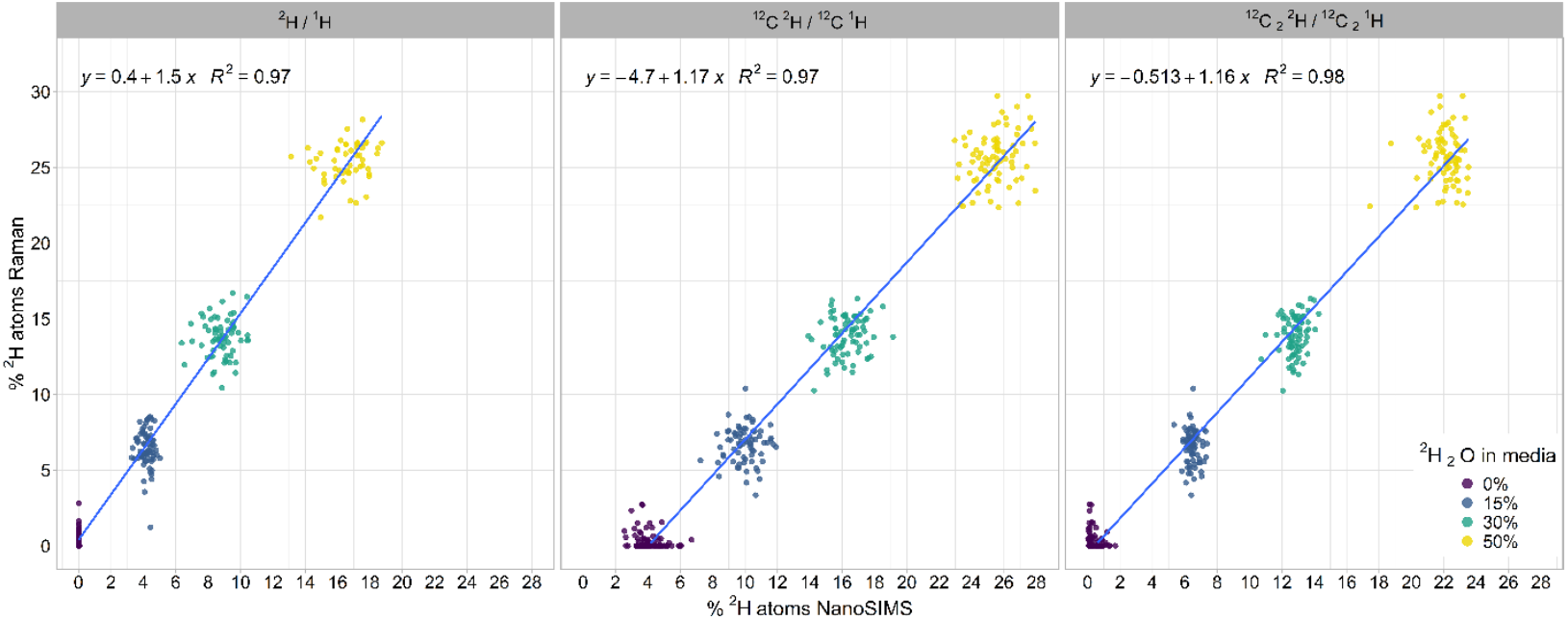
Single cell comparison of the ^2^H content of cells as measured by Raman and NanoSIMS using specific mass ratios. Each dot represents a single cell analyzed with both techniques for different ^2^H_2_O concentrations in the culture medium. The linear model equation and fit (blue line) is shown for each comparison of Raman to NanoSIMS regarding the specific mass ratios.

**Fig. 3.**
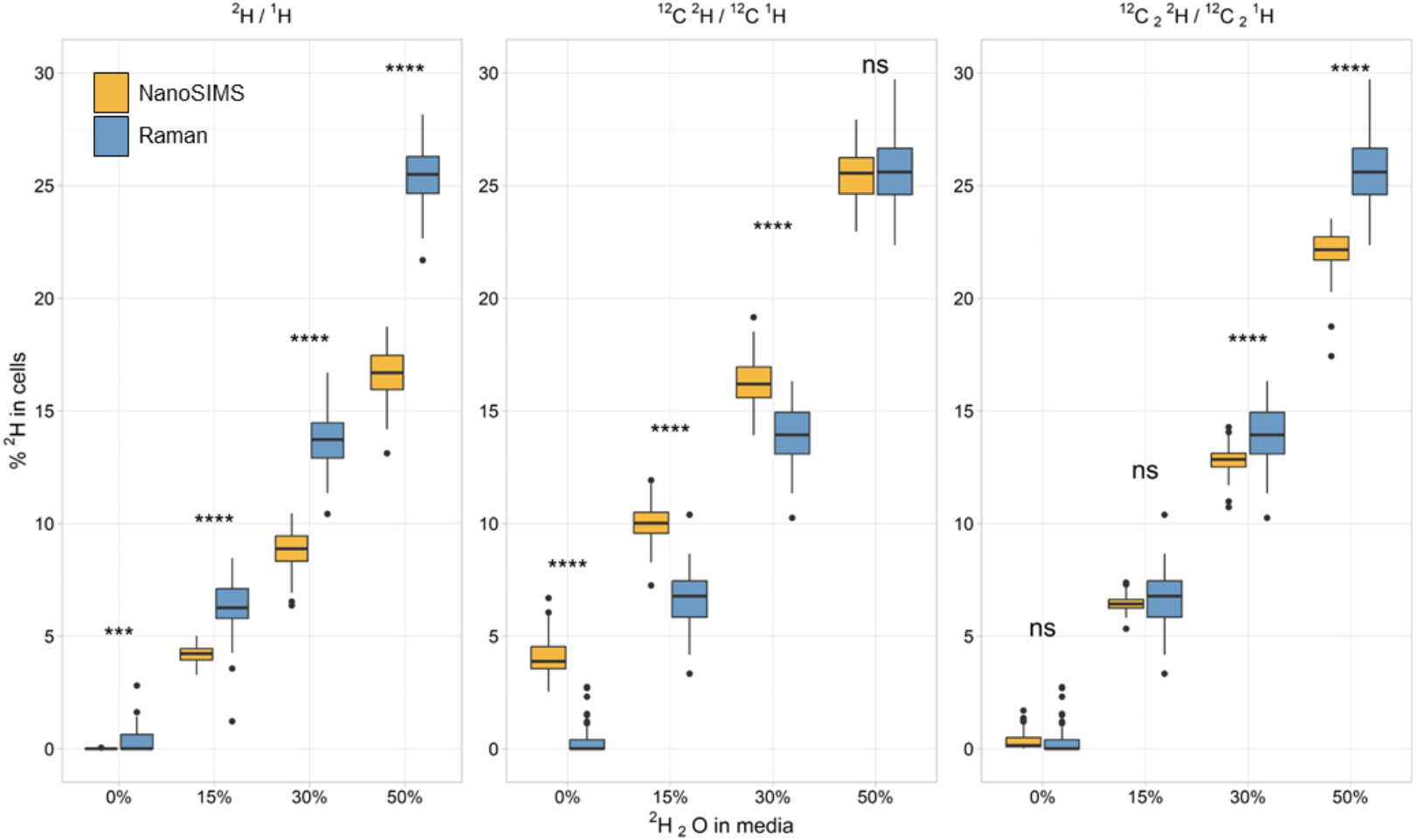
Comparison of ^2^H atom percent as measured by Raman (2040-2300 cm^-1^ wavenumber range for ^2^H peak) and NanoSIMS using the three different mass ratios. A higher ^2^H content was measured by Raman as compared to the NanoSIMS ^2^H/^1^H mass ratio measurements. The opposite is true for the ^12^C^2^H/^12^C^1^H mass ratio for which Raman measures a lower ^2^H content than NanoSIMS. The ^12^C_2_^2^H/^12^C_2_^1^H mass ratio had more comparable results to Raman for the 0% and 15% ^2^H_2_O incubations but measured a lower ^2^H content than Raman for the 30% and 50% incubations. All statistically significant differences are shown: *** = p-value < 1.0 × 10^−3^, **** = p-value < 1.0 x10^−10^, ns = not significant.

### Analysis of Raman wavenumber range

The wavenumber range of 2,040-2,300 cm^-1^ for Raman analysis of ^2^H incorporation was originally proposed by Berry *et al*. and has since been ubiquitously applied in similar studies (2, 16). However, we observed that applying this range uniformly to calculate ^2^H content in cells with lower incorporation levels, such as those incubated in 15% ^2^H_2_O, could result in baseline values being incorporated in final deuterium content (Fig 1A and S3). This discrepancy could affect the calculated ^2^H values. Furthermore, the exact destination of ^2^H within a cell is dependent on the metabolism of the organism (*e*.*g*. anaerobic/aerobic and autotrophic/heterotrophic) and specific wavenumber peak can be influenced by additional available substrates in the incubation, highlighting the nuances to individual datasets (2, 52). To address this, we investigated whether an empirically derived wavenumber range would yield results comparable to those obtained by NanoSIMS.

To assess the potential for alternative wavenumber ranges, we used the 128 Raman spectra collected from cells grown in 50% ^2^H_2_O. These spectra were processed using Savitzky-Golay smoothing, SNIP baselining, and sum normalization. The purpose of using the 50% ^2^H_2_O data was two-fold. First, the higher ^2^H_2_O concentration produces a more normal distribution of the ^2^H peak, thereby reducing the noise that may influence the processing. Second, it ensured the inclusion of all relevant data since lower ^2^H_2_O concentrations would constrain the wavenumber range and likely result in removal of data at higher concentrations. Raman spectra were truncated to 2,000-2,400 cm^-1^ and further smoothed using a spline function (53). Spline smoothing is a nonparametric regression technique used to estimate a curve directly from noisy data without relying on specific parametric models, thus reducing the impact of inherent noise in the spectra on the wavenumber range calculations. The smoothness of the curve is influenced by the degrees of freedom (DOF) of the function, which can dramatically affect the calculation of wavenumber ranges. To evaluate the impact of smoothing on wavenumber ranges, we used a stepwise strategy using various DOF (Fig. S10). We computed the 2^nd^ derivative from the smoothed curve to identify the inflection points (*i*.*e*., where the 2^nd^ derivative changes signs) and used the wavenumber values at which the inflection points occurred to determine the wavenumber range used to calculate the AUC (Fig. S11 and S12). This approach provides a standardized method for determining the wavenumber range used to calculate ^2^H, yielding an average wavenumber value for the DOF of the 128 Raman spectra analyzed (Table S4).

Using the derived wavenumber ranges for each DOF, we calculated the ^2^H atom percent for the 50% ^2^H_2_O Raman data and compared the results to those obtained using the canonical 2,040-2,300 cm^-1^ range and NanoSIMS data (Fig. S13). Notably, the canonical 2,040-2,300 cm^-1^ wavenumber range produced ^2^H values that closely aligned with NanoSIMS ^12^C^2^H/^12^C^1^H mass ratio values, suggesting that historic studies using this wavenumber range and mass ratio have yielded reliable results. Our analysis indicated that a narrower wavenumber range calculated from 10 DOF resulted in ^2^H concentrations that were statistically indistinguishable from the ^2^H/^1^H mass ratio (Table S5). Similarly, no statistical difference was found when comparing the ^2^H atom percent in Raman spectra analyzed using 5.57 DOF to the NanoSIMS ^12^C_2_^2^H/^12^C_2_^1^H mass ratio. When applied to the 0%, 15%, 30%, and 50% ^2^H_2_O datasets, these optimized wavenumbers yielded ^2^H values that were comparable to the NanoSIMS results with lower statistical variance across the datasets of the respective mass ratio (Fig. 4, S13 and Table S6). Linear regressions of individual cell data further supported these findings, showing stronger correlations and improved alignment between the two techniques for all incubation conditions (Fig. S14). As compared to the ^2^H atom percent calculations using the canonical 2,040-2,300 cm^-1^ range shown in Figure 2, wavenumber ranges calculated using the 2^nd^ derivative approach had closer to 1, suggesting a stronger linear correlation between the data when employing wavenumber ranges calculated using the second derivative approach (Table S7). This analysis demonstrates that calculating wavenumber ranges using the second derivative approach improves the consistency and accuracy of Raman-based ^2^H quantification, particularly in studies with varying ^2^H_2_O concentrations.

**Fig. 4.**
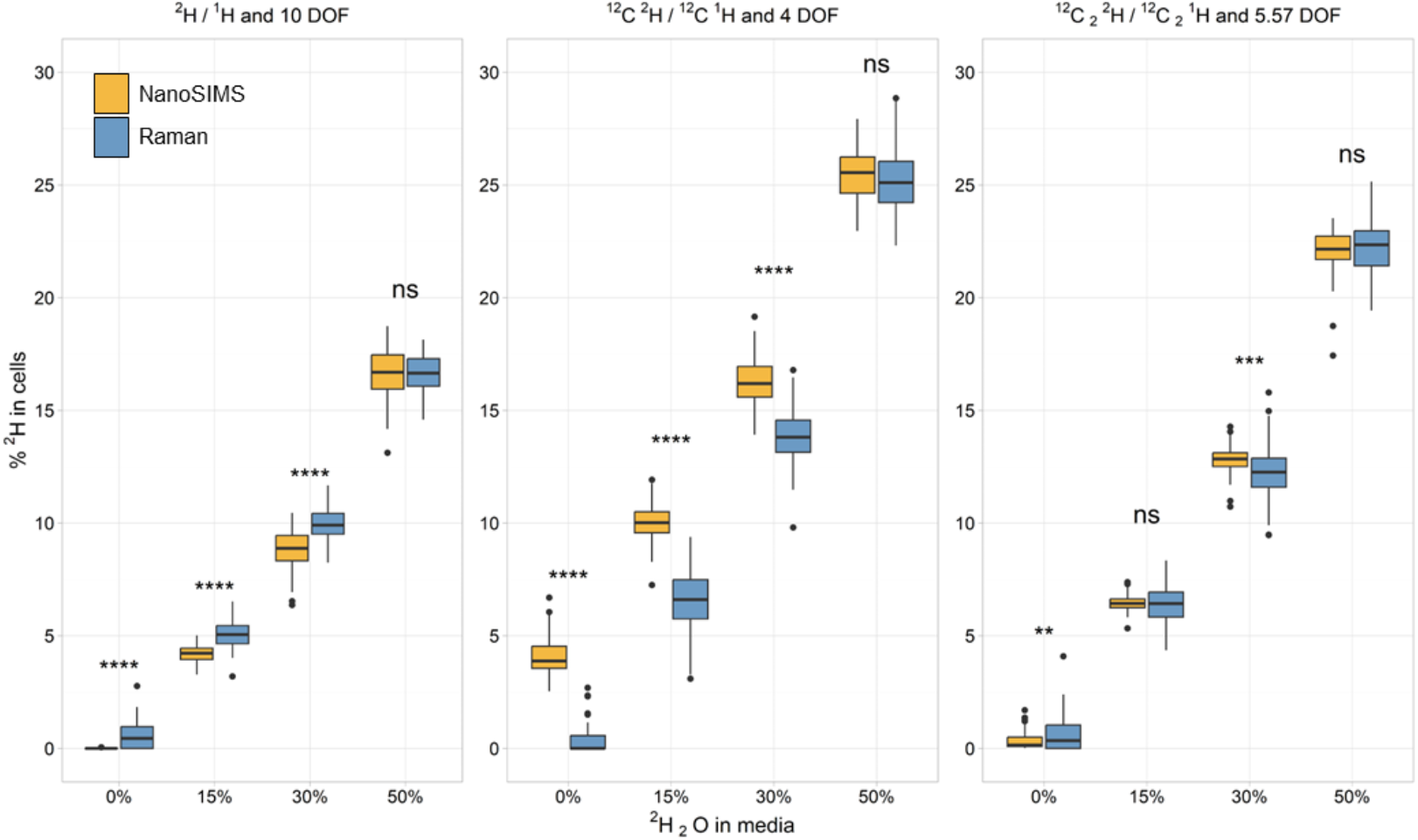
Comparison of ^2^H atom percent as measured by Raman using wavenumber ranges determined by DOF (see Table S4) compared to the three mass ratios measured using NanoSIMS. Using the wavenumber ranges deduced from the 2nd derivatives resulted in Raman data more closely resembling the NanoSIMS measurements for the ^2^H/^1^H and ^12^C_2_^2^H/^12^C_2_^1^H mass ratios. All statistically significant differences are shown: *** = p-value < 1.0 × 10^−3^, **** = p-value < 1.0 x10^−10^, ns = not significant.

## Conclusion

Single species microbial populations have been shown to exhibit metabolic heterogeneity, requiring analysis on a single cell level to differentiate phenotypes within the population (54-56). The application of SIP in conjunction with Raman or NanoSIMS is a common approach to studying nuanced physiological differences in populations (3, 12, 16) and for determining the metabolic activity of cells (14, 18, 57-59). Our work, together with the recent study by Caro *et al*. (26), enables a direct comparison and statistical analysis of Raman and NanoSIMS performed on single-cells using different concentrations of ^2^H_2_O. By quantifying the uptake of ^2^H into the biomass of individual *E. coli* cells using correlative SIP-Raman-NanoSIMS, we established the extent of data equivalence between the two techniques, improving the comparability of studies employing either method.

Several factors were considered before comparing datasets. First, we assessed statistical pre-processing methods for Raman data showing that for *E. coli* K12, truncation and various baselining methods have little impact on the calculation of ^2^H content (Fig. S4 and S5). While pre-processing methods had negligible influence on silent region of the spectra, it is likely that there would be a measurable effect on the fingerprint region of the spectra, highlighting the importance of testing the different pre-processing methods prior to data analysis (31). Additionally, we investigated ^2^H measurement using different masses for NanoSIMS and found that ^2^H measured in cells varies depending on the mass analyzed (Fig. S8). This variation could be due to a combination of different IMF factors, variable atomic mixing during the sputtering process, hydrogen background in the instrument, and sampling of different ^2^H populations pools within the cell (22, 60). For example measurements using ^2^H/^1^H mass ratio include all sputtered ^2^H and ^1^H, whereas ^12^C^2^H/^12^C^1^H and ^12^C_2_^2^H/^12^C_2_^1^H mass ratios specifically sample ^2^H and ^1^H that is bound to ^12^C (22, 61). This difference introduces a selection bias, as these mass ratios exclude hydrogen bound to other atoms, such as nitrogen or oxygen, that are not proportionally measured. Raman measurements exhibit a similar bias, as they detect ^2^H bound to various biomolecules.

Comparison of Raman and NanoSIMS data measuring the anabolic metabolism of ^2^H in single cells demonstrated areas of similarity, supported by robust correlation coefficients for data analyzed using both the canonical Raman wavenumber range for and the 2^nd^ derivative approach (Fig. 2 and S13). Despite these correlations, the inherent methodological differences between Raman and NanoSIMS highlight the need for careful consideration of their respective strengths and limitations. These results support the use of either Raman or NanoSIMS when analyzing cells incubated with ^2^H_2_O, offering researchers flexibility in choosing the most suitable technique for their specific experimental requirements while allowing comparability to literature data.

The decision to use Raman versus NanoSIMS depends on the experimental question. Modern Raman instruments enable faster measurements than NanoSIMS and the infrastructure cost is several times lower. Acquisition times for spontaneous Raman, such as the instrument used in this study, typically range from 1-60 seconds per measurement, but more advanced systems such as Stimulated Raman spectroscopy can measure a field of view, which can contain hundreds of cells, within a minute (16). Raman analysis directly targets C-H bonds, simplifying the interpretation of activity measurements. Confocal Raman microspectroscopy can make point measurements at a resolution of ca. 0.3 μm, making it well suited for analyzing microbial communities (16). For NanoSIMS analysis, ^2^H/^1^H measurements have very low detection limits enabling natural abundance measurements, despite having a relatively low electron affinity (0.75 V; (62)). Assuming three standard deviations for detection limits, the uncalibrated detection limit for the present experiment for NanoSIMS is 0.04 at% compared to >0.2% for Raman. However, NanoSIMS measurement targets all H in the cell including exchangeable H (*e*.*g*., hydroxyl, amino, or thiol groups) as well as residual water. Thus, isotopic exchange of ^2^H during incubation with very minimally active cells could result in ‰ changes that affect the results (26). The ^12^C^2^H/^12^C^1^H ion pair targets C-H bonds directly, but high background, presumably due to ^12^C^1^H_2_, prevents low detection limits at the high-transmission analysis conditions used here. An unknown amount of atomic recombination probably occurs so some dilution of the ^12^C_22_H signal is possible (22). ^12^C_2_^2^H/^12^C_2_^1^H measurement can be performed at higher precision, but again recombination and background interferences limit detection and potentially accuracy. On the positive side, depending on enrichment levels, other peaks could be monitored along with either polyatomic pair to measure assimilation of other elements beside H (44). When lower detection limits than those afforded by ^12^C^2^H/^12^C^1^H or ^12^C_2_^2^H/^12^C_2_^1^H are required or the experiment requires the measurement of all cellular ^2^H, we recommend that researchers consider using ^2^H/^1^H, in addition to other mass ratios.

Our comparative investigation fills a gap in the current understanding of spectroscopy and spectrometry techniques in microbiology and offers a foundation for the further exploration and application of SIP methods in diverse microbial systems. The comparability of Raman and NanoSIMS measurements of single cells presented in this study paves the way for a more nuanced and comprehensive understanding of microbial activities, interactions, and responses to isotopic labeling.

## Materials and Methods

### Preparation of isotopically labeled cells

*Escherichia coli* K12 (DSM498) cultures were grown aerobically at 200 rpm agitation at 37 °C for 13 h in M9 medium (0.38 mM thiamine and 22.2 mM glucose) that had been amended to a final concentration of either 0%, 15%, 30%, or 50% ^2^H_2_O (99.9%-^2^H; Cambridge Isotope Laboratories). The optical density (600 nm) of the cultures was measured every hour for the first six hours, after which measurements were taken every half hour. When the OD_600_ had reached 0.8-0.9, 1 mL of culture was sampled (Fig. S1) and chemically fixed by adding paraformaldehyde (PFA; Electron Microscopy Science; EM grade) to a final concentration of 2%. These samples were then incubated for 60 min at room temperature. Cells were then washed twice with 1× phosphate buffered saline (PBS; pH 7.4) by centrifugation at 16,000 g for 5 min, after which supernatants were removed, and cell pellets were resuspended in 1 mL of 1x PBS and samples stored at 4 ºC.

### Preparation of stainless steel coupon

Fixed cells were dried to the surface of mirrored stainless steel due to its desired properties as a Raman substrate (63). The stainless steel coupon was prepared as previously described in Schaible *et al*. (25). Briefly, the coupon was cleaned by washing with a 1% solution of Tergazyme (Alconox, New York, NY) rinsed with Milli-Q water, subsequent one-minute washes in acetone and 200 proof ethanol, and then air dried. To maintain correct orientation of the samples, asymmetric boxes were etched into the mirrored surface of each coupon using a welder’s pen (Fig. S2). 1 μL of each sample was spotted onto the coupon and air-dried at 46 °C for 1 min, after which the coupon was washed in 1:1 ethanol:MilliQ for 1 minute and then air dried. The coupon was incubated in a 300 nM (100 ng/mL) solution of DAPI (4’,6-diamidino-2-phenylindole, ThermoFisher, Waltham, MA) for 3 minutes as per the manufacturer’s instructions. The coupon was then rinsed three times in fresh PBS and then briefly dipped them into ice-cold Milli-Q water to remove salts and air-dried using compressed air. To track single cells across platforms, regions of interest (ROIs) were drawn around cells on the coupon using a Leica LMD6 Laser Microdissection System (Danaher Corporation, Washington DC) using a power setting of 7, a speed of 7, and aperture size of 7 (Fig. S3).

### Confocal Raman microspectroscopy and spectral processing

Raman spectra of individual cells were acquired using a LabRAM HR Evolution Confocal Raman microscope (Horiba Jobin-Yvon, located at Montana State University in Bozeman, Montana) equipped with a 532 nm laser and 300 grooves/mm diffraction grating. The instrument was calibrated daily before use with a silica oxide standard. Cell spectra were acquired using a 100× dry objective (NA = 0.90) in the range of 250–3200 cm^−1^, with 3 acquisitions of 10 s each, and a laser power of 4.5 mW. Spectra were processed using LabSpec version 6.5.1.24 (Horiba). The spectra were preprocessed in GNU R (64) using the Alkahest (65), ptw (66), and Pracma (67) packages. Each spectra was smoothed using the Savitsky-Goley algorithm, (34) followed by background subtraction and baselining of spectra using either asymmetric least squares (ALS) (35), polynomial fit (36), rolling ball (37), or statistics-sensitive non-linear iterative peak-clipping (SNIP) (38). Raman spectra baselined with the SNIP method were used for the comparison to the NanoSIMS data. Settings used for smoothing and baselining can be found in the R file deposited on GitHub (https://github.com/georgeschaible/Raman-spectra-processing). Each spectrum was then normalized to the sum of its absolute spectral intensity, and the incorporation of ^2^H into biomass was calculated by 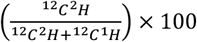, using the integration of the area under the curve (AUC) for the ^12^C^1^H (2800–3100 cm^-1^) peak, the canonical ^12^C^2^H (2040–2300 cm^-1^), or the wavenumber ranges calculated using the 2^nd^ derivative (Table S4). The AUC was calculated using the trapezoidal rule as follows: 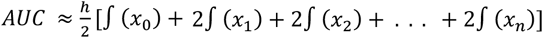where *h* is the width of each interval, ∫*x*_*i*_ is the function value at each data point, and *n* is the number of data points.

### Nano-scale secondary ion mass spectrometry

To map the elemental composition and the relative isotopic abundances of hydrogen, ion images of the same individual cells investigated by Raman were acquired using the Cameca 50L NanoSIMS at the Environmental Molecular Sciences Laboratory at Pacific Northwest National Laboratory in Richland, Washington. The vacuum gauge pressure in the analytical chamber during all analyses was consistently less than 3 × 10^−10^ mbar. Samples were stored in a dry N_2_ storage box when not under instrument vacuum. Although no isotopic standards were available to calibrate the instrument for instrumental mass fractionation (IMF) similar responses for different multipliers were ensured by setting electron multiplier (EM) voltages such that the maximum pulse height distribution (PHD) was at a discriminator voltage of - 230 ± 10 mV. ^1^H was used to set the PHDs for ^2^H^-^/H^-^ analyses and ^28^Si^-^ was used to set PHDs for ^12^C_2_^1^H^-^/^12^C^1^H^-^ and ^12^C_22_H^-^/^12^C_2_^1^H^-^ measurements. For ^12^C_2_^1^H^-^/^12^C^1^H^-^ and ^12^C_22_H^-^/^12^C_2_^1^H^-^ analyses, conditions included a 200 μm D1 aperture, 20 μm entrance slit, 200 μm aperture slit, and 90 μm exit slits. ^2^H^-^/^1^H^-^ analyses used a 200 μm D1 aperture, 30 μm entrance slit, 350 μm aperture slit, and 90 μm exit slits. All NanoSIMS images were acquired using a 2 pA (∼120 nm) 16 keV Cs^+^ primary ion beam at 256 × 256-pixel resolution with a dwell time of 1 ms px^−1^. Analysis areas were pre-sputtered with ∼5 × 10^15^ ions cm^−2^ prior to collecting 15 consecutive frames. Secondary ions (^12^C^1^H^-, 12^C^2^H^-, 12^C_2-_, ^12^C_2_^1^H^-, 12^C_22_H^-, 35^Cl^-^, and ^12^C^14^N^16^O^-^ or ^1^H^-^ and ^2^H^-^) were accelerated to 8 keV and counted simultaneously using the EMs. ^35^Cl^-^, and ^12^C^14^N^16^O^-^ were not used for further analyses. Detectors collecting ^2^H^-^ and ^1^H^-^ ions were situated near the center of the magnet radius to improve simultaneous secondary centering characteristics, with careful attention paid to the secondary tuning of the Helmholtz coils to align ^2^H and ^1^H. Because different tunning is required for simultaneous ^12^C^2^H^-^/^12^C^1^H^-^ and ^12^C_22_H^-^/^12^C_2_^1^H^-^ compared to ^2^H^-^/^1^H^-^, these measurements were performed on different cells. After pre-sputtering, secondary signal was centered prior to collecting 15 consecutive frames for each image. The OpenMIMS plugin for ImageJ was used to correct images pixel by pixel, for dead time (44 ns) and Quasi Simultaneous Arrivals (QSA; β = 0.5), prior to aligning and summing the 15 frames. There is a small amount of H in the stainless steel wafers that we used for the analyses. We found that ^2^H^-^/^1^H data were sensitive to ROI size with respect to the *E. coli* ion image size. Therefore, to optimize reproducibility, ROIs, of single microbes were acquired using the Bemsen algorithm in the Auto Local Threshold function of ImageJ on ^12^C_2_^-^ or ^1^H aligned and summed images. Examples of ROIs are shown in Figure S15. Similar external and average internal uncertainties within treatments for the ^2^H/^1^H measurements indicate that the degree of H interference from the stainless steel substrate was minimal (Table S8). Data from single microbe ROIs were exported to a custom spreadsheet for data reduction. Internal uncertainties were obtained from counting statistics (68).

### Preparation of fatty acid methyl esters

The *E. coli* sample was freeze-dried overnight. Preparation of fatty acid methyl esters (FAMEs) was done following the procedure adapted from Rodriguez-Ruiz et al. (28), discussed briefly here. 2 mL of a mixture of 20:1 v/v anhydrous methanol/acetyl chloride and 1 mL of hexane were added to 1.3 mg of the sample in a glass culture tube. A method blank (culture tube with no sample) was also included to investigate whether there were any contaminations resulting from the method. The two glass tubes were capped tightly and heated at 100 °C for 10 min and then allowed to cool to room temperature. 2 mL of deionized water and 2 mL hexane added forming two phases. The hexane (top) phase was removed and collected in a 40 mL VOA vial. The extraction was repeated with further 2 mL additions of hexane to maximize the extraction yield. The sample and the blank were concentrated to 0.125 mL under nitrogen gas. The FAMEs peaks present in the sample and blank were identified via gas chromatography/mass spectrometry (GC/MS) on a ThermoScientific Trace 1310 TSQ-9000 equipped with a Zebron ZB-XLB column (30 m x 0.25 mm I.D. x 0.25 μm film thickness) and programmable temperature vaporizing (PTV) injector operated in split mode (split ratio 41.7), using helium as a carrier gas (flow rate = 1.2 ml/min). The GC oven was held at 100 °C for 1.5 min, ramped at 20 °C/min to 150 °C with no hold, then ramped at 5 °C/min to 300 °C with a final 2 min temperature hold. Peaks were identified by comparing the relative retention times and mass spectra to those an eight-compound FAMEs standard mixture, as well as to mass spectra in the NIST MS Library database.

### δ^2^H isotope analysis

The δ^2^H values of the FAMEs in the sample were measured five times by a gas chromatograph coupled to an isotope ratio mass spectrometer (Thermo Scientific 253 Plus) using a pyrolysis interface (*i*.*e*., GC/P/IRMS). Chromatographic separation was achieved on a thick-film Agilent DB-XLB column (30 m x 0.25 mm I.D. x 1.00 μm film thickness) and PTV injector operated in splitless mode, using helium as a carrier gas (flow rate = 1.4 ml/min). The GC oven was held at 120 °C for 1 min, ramped at 120 °C/min to 225 °C with no hold, ramped at 0.5 °C/min to 240 °C, and then ramped at 10 °C/min to 325 °C with a final 5 min temperature hold. Measured isotope ratios were calibrated using hydrogen gas of known isotopic composition and are reported in δ notation (in units of ‰, or parts per thousand) relative to the Vienna Standard Mean Ocean Water (VSMOW) international standard (δ^2^H = RAA/RVSMOW − 1), where R = ^2^H/^1^H. Additionally, a FAME standard was analyzed to verify instrument accuracy and precision.

### Statistical analyses

All datasets were analyzed in GNU R (64) using the tidyverse, rstatix, emmeans, and ggpubr packages (69-71). Statistical differences between multiple variables were determined using pairwise t-tests with a Bonferroni p-adjusted method. Calculations to determine the wavenumber range for ^2^H used the pspline R package (72) and in-house written R code for the calculation of 2^nd^ derivative inflection points of smoothed curves. Code used for analysis is deposited on GitHub (https://github.com/georgeschaible/Raman-spectra-processingd).

## Data availability

Raman spectra have been deposited in the MicrobioRaman database under accession number S-MBRS8. NanoSIMS raw data is stored long-term on the EMSL data repository (search.emsl).

## Author contributions

GS and RH designed the study. GS performed culturing and prepared samples for analysis. JAC collected the Raman spectra and GS analyzed the spectra. JBC and JB performed NanoSIMS analyses. GS and JBC compiled the data. MM performed the lipid extraction and esterification, conducted the IR/MS analysis, and collaborated with AS to analyze the data. JA and GS performed the second derivative calculations. GS performed statistical analyses and prepared figures. GS, JBC, and RH wrote the manuscript draft, which was then edited by all authors.

## Acknowledgements

This study was funded through NIH award 1R35GM147166-01 to R.H. GS was partially supported by NASA FINESST fellowship 80NSSC20K1365. JAC was supported by NSF Research Traineeship Program award 2125748. Montana State University’s Confocal Raman microscope was acquired with support by the NSF MRI program (DBI-1726561) and the M. J. Murdock Charitable Trust (SR-2017331). A portion of this research was performed under the Facilities Integrating Collaborations for User Science (FICUS) program (awards DOI: 10.46936/fics.proj.2020.51544/60000211) and used resources at the Environmental Molecular Sciences Laboratory (https://ror.org/04rc0xn13), which is a DOE Office of Science User Facilities operated under Contract No. DE-AC05-76RL01830. This manuscript has been authored by UT-Battelle, LLC, under contract DE-AC05-00OR22725 with the US Department of Energy (DOE). The US government retains and the publisher, by accepting the article for publication, acknowledges that the US government retains a nonexclusive, paid-up, irrevocable, worldwide license to publish or reproduce the published form of this manuscript, or allow others to do so, for US government purposes. DOE will provide public access to these results of federally sponsored research in accordance with the DOE Public Access Plan (https://www.energy.gov/doe-public-access-plan). We thank Hope McWilliams (MSU) for assistance with SIP labeling and Anthony Kohtz (MSU) for helpful input on the analysis of Raman spectra.

## Notes

**Conflict of interest:** The authors declare no conflict of interest.

### Competing Interest Statement

The authors have declared no competing interest.

### Summary of Updates

revised version of the study that includes new IRMS measurements of cultures incubated without heavy water; revised results and discussion

