## Supplementary material for "Comparing Raman and NanoSIMS for heavy water labeling of single cells": SI text

### NanoSIMS analysis: methodology, considerations, and sources of variability

Because we wanted to perform spatially resolved isotopic analyses on single microbes, we chose to use a low primary beam current with high spatial resolution (2pA, ~120 nm). The requirement for spatially resolved single cell analyses precluded us from tuning for high mass resolving power as that would have reduced secondary count rate to unacceptable levels to achieve a robust data set in a reasonable amount of time. Thus, we chose to test the strengths and weaknesses of the various ion pairs when operating at more standard mass resolving power (MRP) conditions. Theoretical MRP for ion species of interest are presented in Table S9 We optimized MRP using the $^{1}H^{-}$, or $^{12}C_2^{-}$ signal from an in-house yeast reference material, MRP for $^{12}C_2^{-}$ was tuned to be greater than 6000 (Cameca definition). For $^{2}H/^{1}H$ analyses, MRP for $^{1}H^{-}$ was greater than 2,000. We emphasize that although we used MRP to tune the instrument for acceptable performance, abundance sensitivity and relative intensity of isobaric interference is more relevant to the amount of interference from adjacent isobaric peaks than MRP. Both metrics are impossible to measure in a meaningful way in many instances during NanoSIMS analyses of complex matrices such as bacteria. Detector deflector values for $^{2}H^{-}$, $C_2H^{-}$, and $C_2{}^{2}H^{-}$ were set to minimize the effect of isobaric interferences using cells grown in 50% $^{2}H_2O$ and were checked at least daily. Examples of deflector tunings for m/z 14 and m/z 26 are presented in Figure S15. It should be emphasized that having fully resolved peaks is not always a requirement for meaningful data acquisition. This is common practice in NanoSIMS analysis protocols as abundance sensitivity rather than MRP is often the more important metric for allowable peak overlap. Common examples often encountered in SIMS analyses are when resolving $^{13}C^{-}$ from $^{12}C^{1}H^{-}$ and $^{12}C^{15}N^{-}$ from $^{13}C^{14}N^{-}$.

Although there was considerable scatter in the NanoSIMS data for all three ion pairs there was clear correlation ($R^2 \geq 0.98$) between $^{2}H_2O$ present in the growth medium and the single cell NanoSIMS signal for all three ion pairs (Fig. 3 and S6). Slopes for regressions between $^{2}H_2O$ in the growth medium and $^{12}C^{2}H^{-}/^{12}C^{1}H^{-}$, and $^{12}C_2{}^{2}H^{-}/^{12}C_2{}^{1}H^{-}$ were not significantly different, but the slope for $^{2}H/^{1}H$ vs growth medium was significantly lower than those of the organic ions. Because we did not have relevant $^{2}H/^{1}H$ standards, we can only speculate on the significance of these slopes, but the equivalent slopes for the $^{12}C^{2}H^{-}/^{12}C^{1}H^{-}$, and $^{12}C_2{}^{2}H^{-}/^{12}C_2{}^{1}H^{-}$ pairs suggest that they are interrogating similar H pool.

Likely contributors to the NanoSIMS data scatter are real variations in cell to cell $^{2}H$ incorporation, variation in the cellular material being analyzed, analytical variability due largely to limited ion counts (counting statistics), variations due to instrument instabilities (due to for example variations in laboratory temperature), and variable contributions due to interferences that have not been fully resolved. Nevertheless, the robust correlations clearly indicate that these measurements have physical meaning. We chose not to subtract the contribution of $^{13}C^{1}H^{-}$ from $^{12}C^{2}H^{-}$ as we set the detection deflector to the middle of the peak which should have minimized abundance sensitivity issues from $^{13}C^{1}H$ (Fig. S16). This, along with the difficult to resolve interference, $^{12}C^{1}H_2^{-}$ are likely responsible for the offset in data from all treatments and positive intercept for this dataset. Because NanoSIMS internal precision is ultimately limited by total ion counts (1), it is interesting to note that, on average, $^{12}C_2{}^{1}H$ counts were roughly five times that of $^{12}C^{1}H$ counts for individual cells (Fig. S6). This is at least in part due to the greater electron affinity of $C_2H$ compared with CH (2, 3). Thus, in theory, the minimum average internal uncertainty (based on counting statistics; (4)) should be roughly 2.2 times higher for $^{12}C^{2}H^{-}/^{12}C^{1}H^{-}$ than for $^{12}C_2{}^{2}H^{-}/^{12}C_2{}^{1}H^{-}$ (4). Table S10 summarizes these data. Except for cells grown without added $^{2}H_2O$, data are largely consistent with these expectations. Cells grown in 15%, 30%, and 50% $^{2}H_2O$ had an average internal standard deviation (SD) of 2.28 for $C^{2}H/C^{1}H$ estimates relative to $C_2{}^{2}H/C_2{}^{1}H$ (n = 230). The corresponding comparison of average external uncertainties is slightly lower (SD = 1.92) and are consistent with expectations of external uncertainties being limited by internal uncertainties. The average relative internal and external uncertainties for the same ion pairs for cells grown without $^{2}H_2O$ are 9.60 and 11.83 respectively (n = 86, Table S10). We attribute the higher uncertainties to the presence of difficult to resolve and highly variable isobaric interferences (1).

Despite our inability to rigorously quantify $^{2}H$ content in *E. coli*, all three ion pairs provided meaningful qualitative data concerning cell activity as measured by $^{2}H$ incorporation. Choice of what ion pair to use may depend on sample quality, degree of $^{2}H$ enrichment, and whether another isotope (e.g. $^{13}C$ or $^{15}N$) has been used as a tracer. Surprisingly, the $^{12}C_2{}^{2}H^{-}/^{12}C_2{}^{1}H^{-}$ ion pair provided high quality data for these analyses and was less affected by natural abundance $^{13}C$ and $^{1}H$ adduct interferences than the $^{12}C^{2}H^{-}/^{12}C^{1}H^{-}$ ion pair. This result occurred even though the m/z peak was difficult to tune to optimize the $^{12}C_2{}^{2}H^{-}$ peak shoulder. Our choice of flat substrate and low instrument vacuum likely eased this burden somewhat and environmental samples with high $H_2O$ content and rough surface topography with variable charging may make this ion pair unusable. Nevertheless, if the ion pair is available from an analytical standpoint, it may provide superior detection limits and count rates compared with $^{12}C_2H^{-}/^{12}C^{1}H^{-}$as it did in this study. Both $^{12}C^{2}H^{-}/^{12}C^{1}H^{-}$ and $^{12}C_2{}^{2}H^{-}/^{12}C_2{}^{1}H^{-}$ would likely need rigorous testing to evaluate their usefulness if large fractions of either $^{13}C$ and/or $^{15}N$ were added as a second tracer. In theory, The NanoSIMS 50L can monitor $^{1}H^{-}$, $^{2}H^{-}$, $^{12}C^{1}H^{-}$, and $^{12}C^{2}H^{-}$ simultaneously (1). The same authors modified the multicollection configuration of their NanoSIMS which is an invasive option that most users likely will like to avoid. Thus far we have found that secondary tuning for $^{2}H^{-}/^{1}H^{-}$ is very different compared with the organic ion pairs for standard NanoSIMS configurations and cannot be coupled with the monitoring of other ion signals if maximum precision is to be maintained. The $^{2}H^{-}/^{1}H^{-}$ provided good ion yields and superior detection limits compared with the organic ion pairs and may be the ion of choice if high accuracy and/or low detection limits are required or, for example, if high levels of $^{13}C$ are used as a secondary tracer. However, this option more than doubles the analysis time required to monitor multiple isotopes.
