## Supplementary material for "Comparing Raman and NanoSIMS for heavy water labeling of single cells": SI figures 1-16

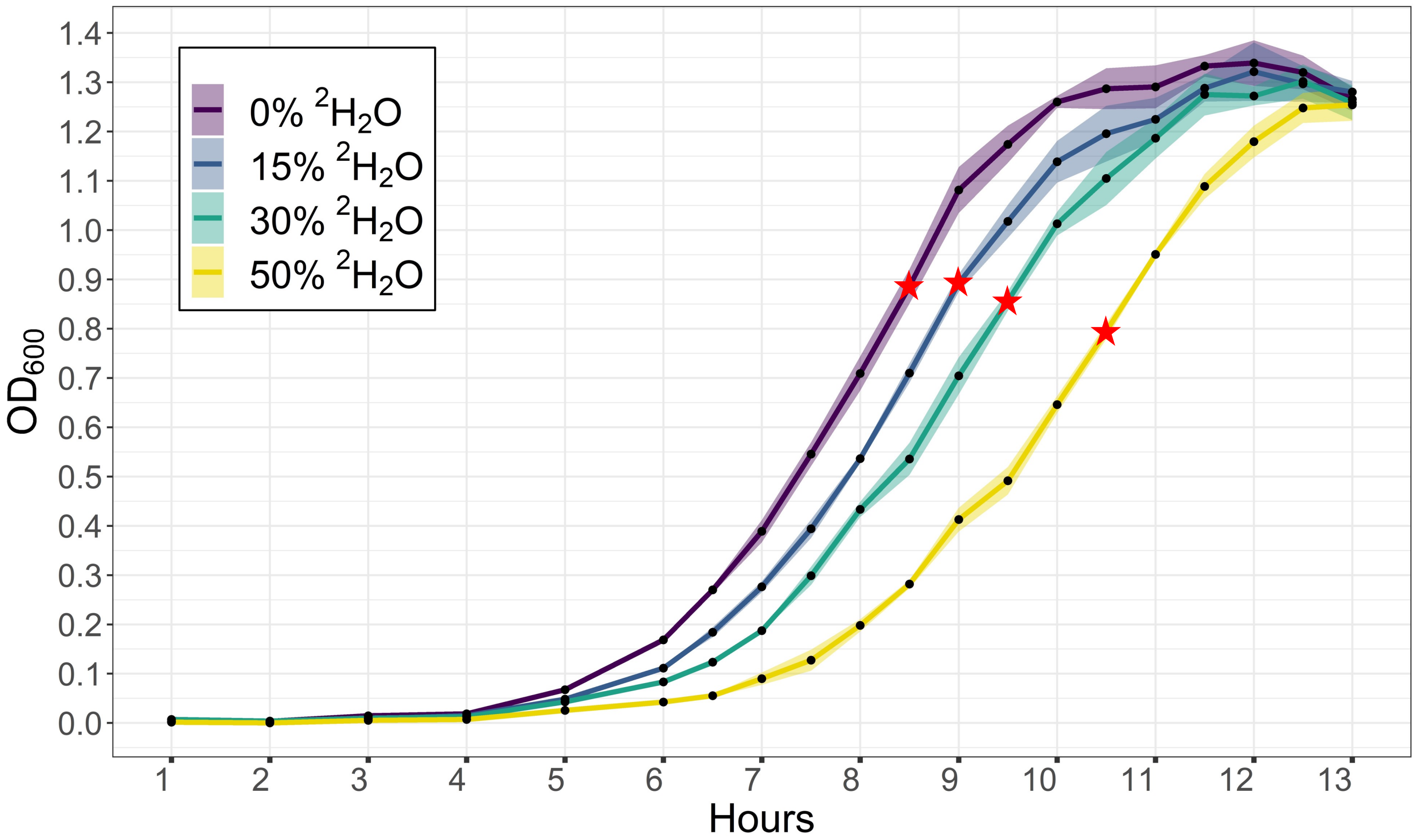


Fig. S1. Growth curves of triplicate *E. coli* cultures grown with varying $^2H_2O$ concentrations. Shaded region behind line shows the minimum and maximum $OD_{600}$ of triplicate samples. Red stars highlight the time and $OD_{600}$ at which cells were sampled and fixed for Raman and NanoSIMS analysis (harvesting at approximately same OD was attempted). The effect of $^2H_2O$ on *E. coli* growth can be seen by a delayed logarithmic growth upon the increased addition of heavy water to the growth media.

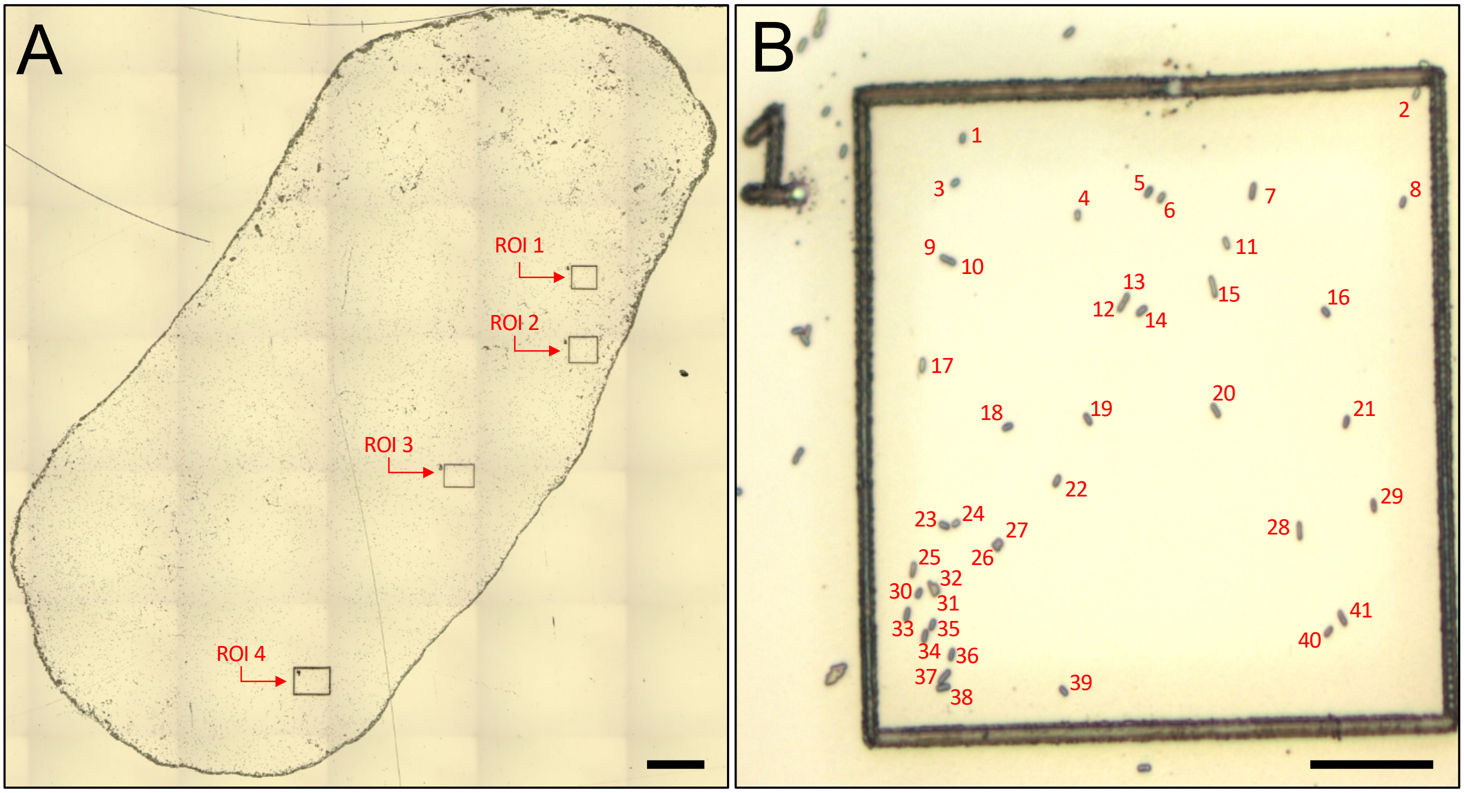


Fig. S2. Example of the mapping of ROIs and cells for correlative workflow. **A** Shows a tiled brightfield microscopy image of the individual ROIs etched into the surface of the stainless steel coupon using a laser microdissection microscope. Scale bar 100 µm. **B** Catalogued *E. coli* cells located within ROI 1 shown in A. Scale bar 10 µm.

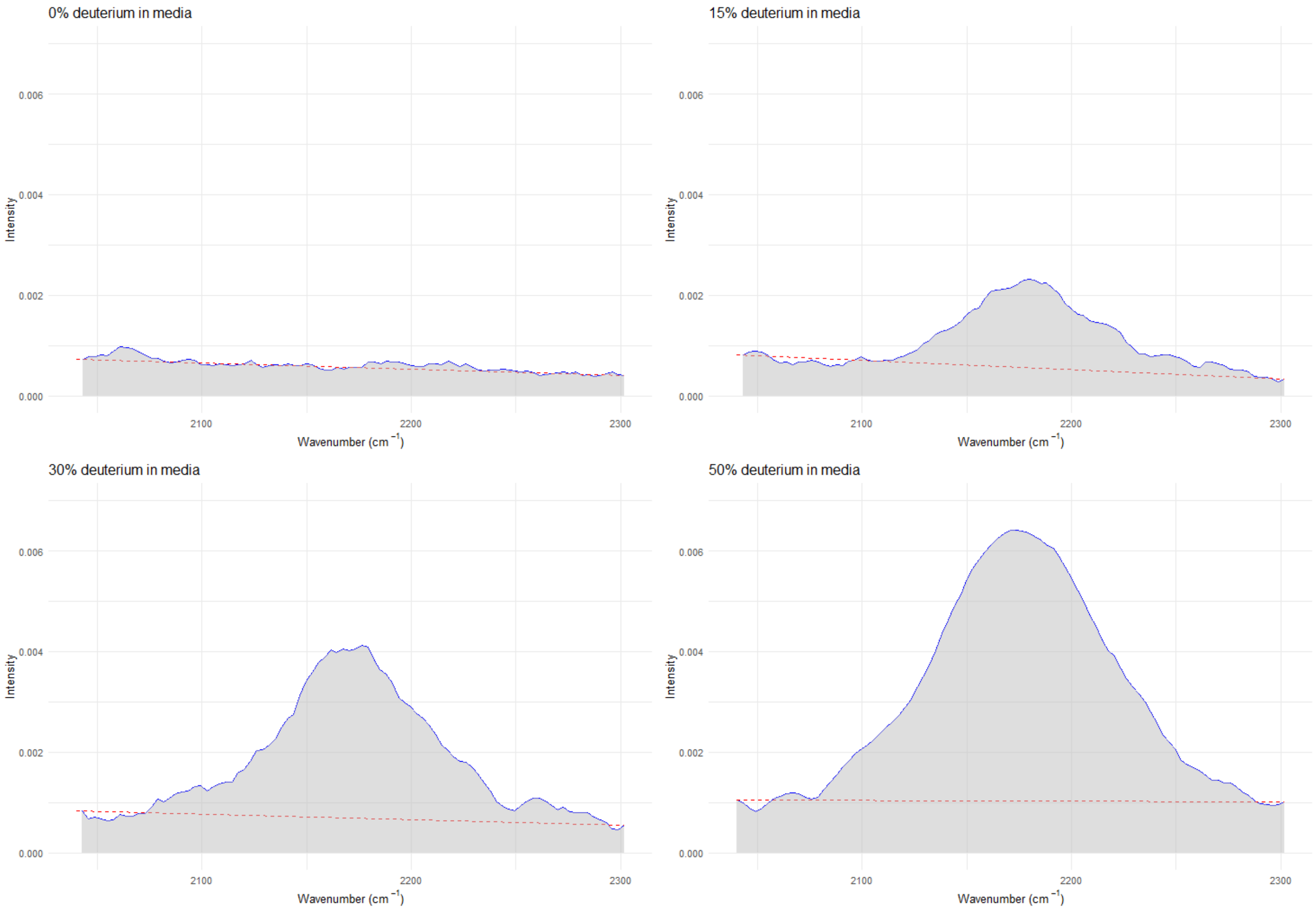


Fig. S3. Example showing the calculation of the AUC for $^{12}C$-$^{2}H$. A baseline is drawn from the starting wavenumber (2,040 cm$^{-1}$) to the ending wavenumber (2,300 cm$^{-1}$) and only the area above that line and below the curve is calculated. The same method was applied to the $^{12}C$-$^{1}H$ peak (2,800-3,100 cm$^{-1}$).

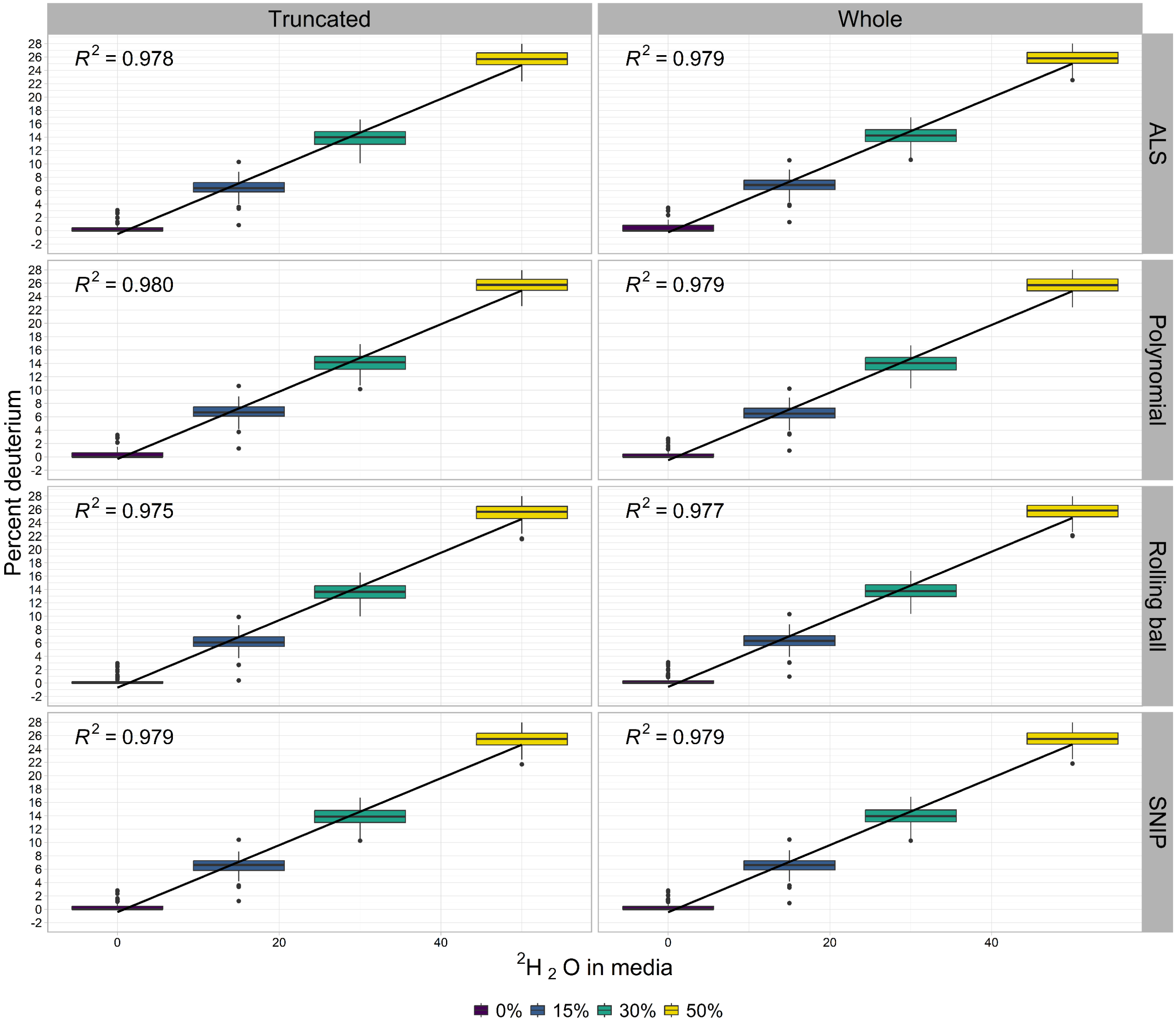


Fig. S4. Analysis of baselining methods for whole and truncated Raman spectra. The percent deuterium was calculated by comparing the ratio between CD and CH of the vibrational bands. Spectra were either analyzed as a whole (250-3,200 $cm^{-1}$) or truncated (1,800-3,200 $cm^{-1}$) with each of the four baselining methods. The analysis showed minimal differences between the baselining methods regarding the r-squared value for each of the concentrations of $^2H_2O$ in the media.

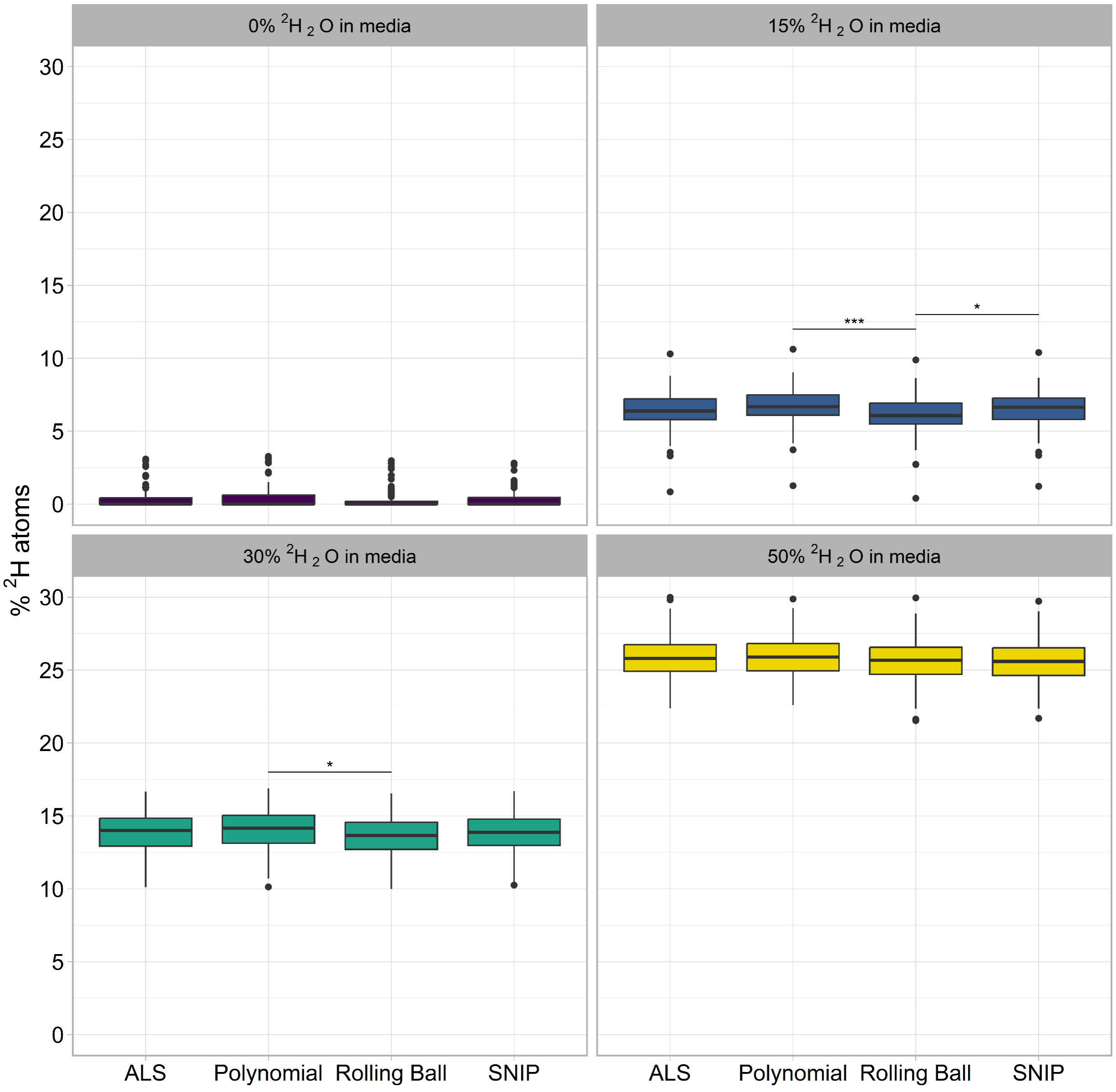


Fig. S5. Analysis of baselining methods for truncated Raman spectra. The percent deuterium was calculated by comparing the ratio between CD and CH of the vibrational bands. Spectra were either analyzed as a whole (250-3,200 $cm^{-1}$) or truncated (1,800-3,200 $cm^{-1}$) with each of the four baselining methods. Only rolling ball baselining method was statistically different as compared to other baselining methods at the 15% and 30% $^{2}H_2O$ incubations.

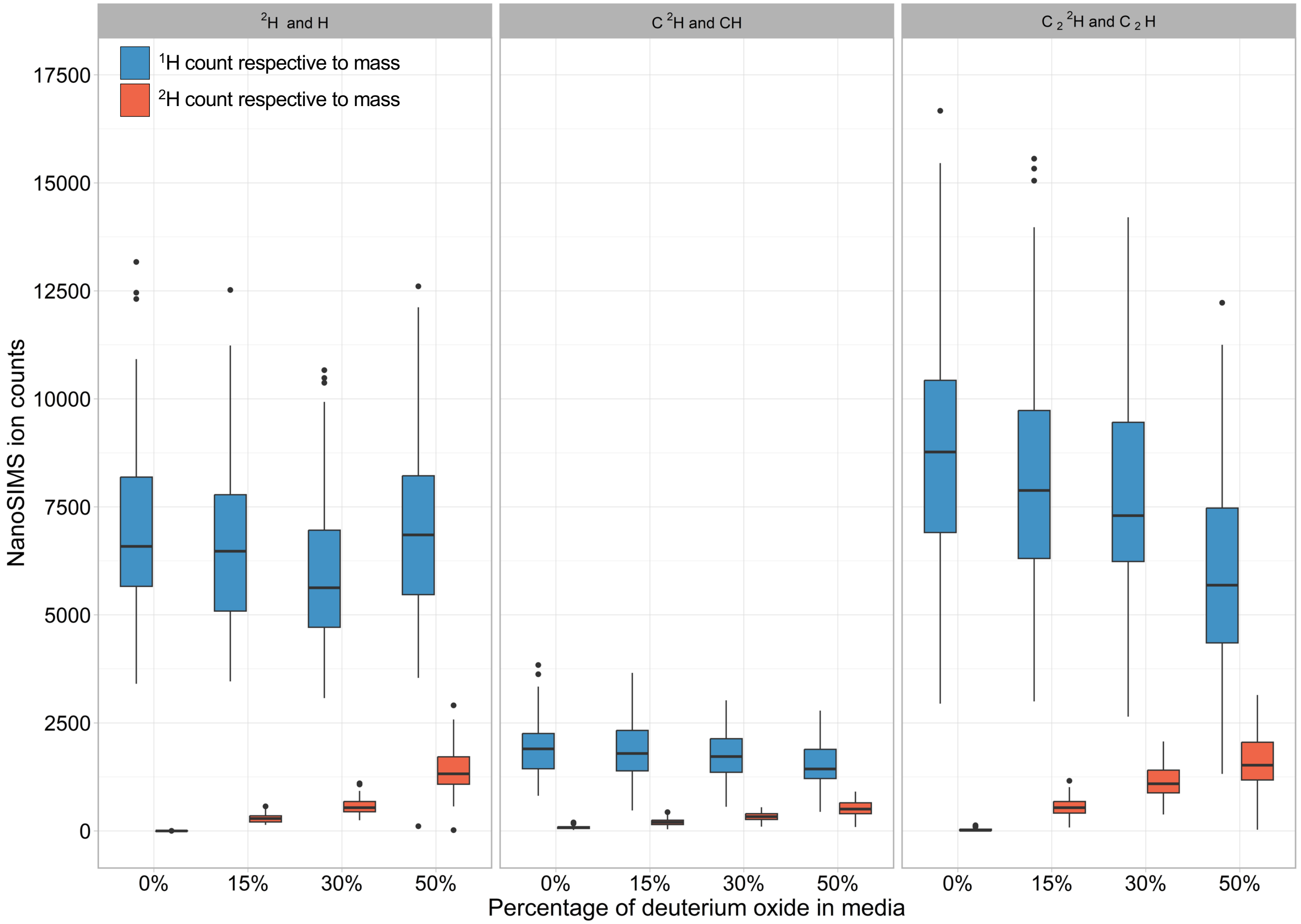


Fig. S6. Comparison of NanoSIMS ion counts for each ion mass analyzed in this study. Blue bars show the mass measured for the H count and orange bars show the mass measured for the $^2$H count, both respective to the mass ratio measured. Both $C^2H/C^1H$ and $C_2{}^2H/C_2{}^1H$ were measured on the same cells, indicating that differences in counts are likely due to instrument drift or kinetic isotope/electron affinity effects.

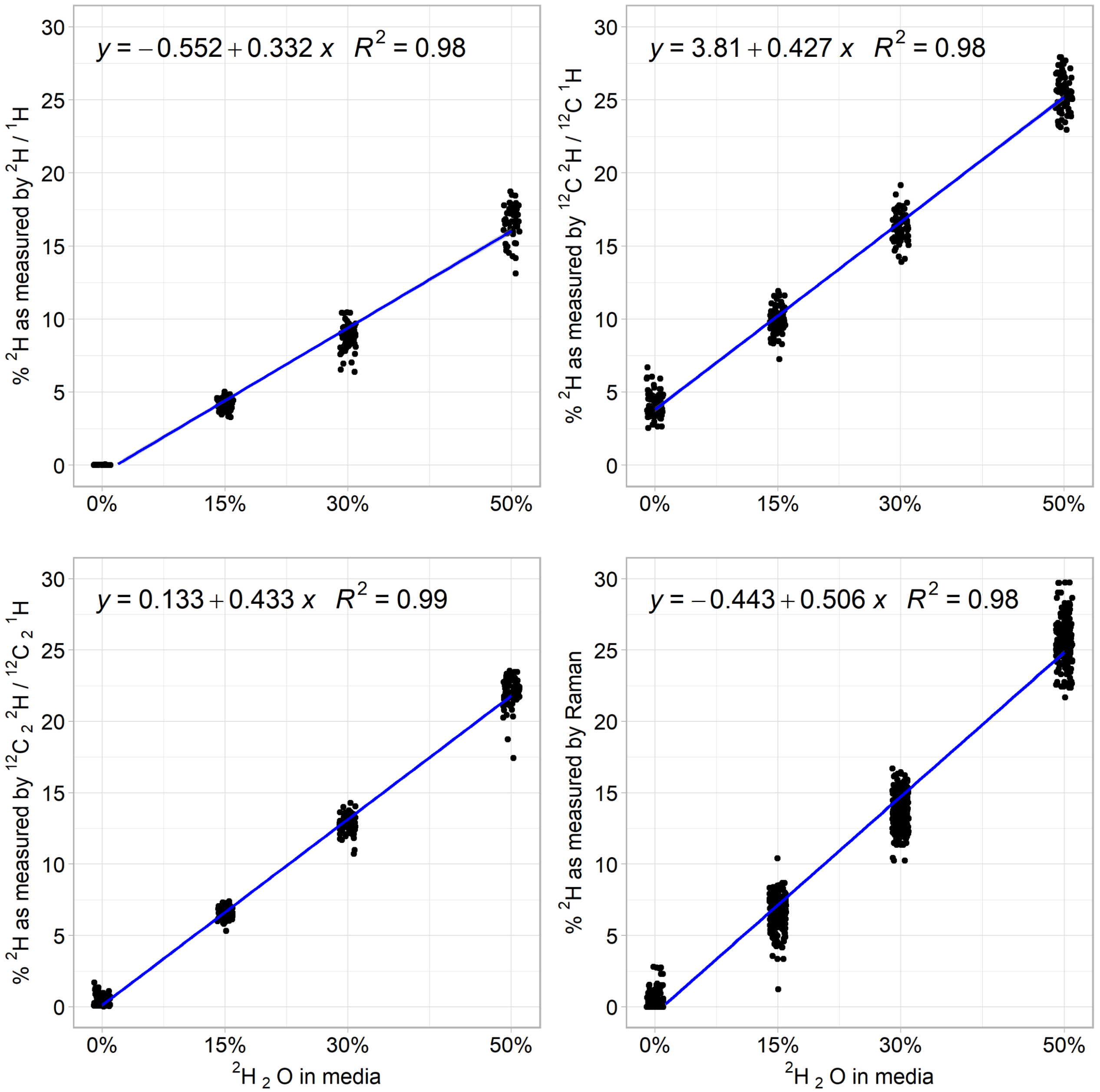


Fig. S7. Linear regressions showing the difference in slope for the $^{2}H$ content of cells measured using the three NanoSIMS masses and by Raman. Both $^{12}C^{2}H/^{12}C^{1}H$ and $^{12}C_2{}^{2}H/^{12}C_2{}^{1}H$ data was collected concurrently and have similar slopes. The $^{2}H/^{1}H$ was collected during a separate analytical session due to the need to adjust detectors for collection and thus cannot be directly compared to the $^{12}C^{2}H/^{12}C^{1}H$ and $^{12}C_2{}^{2}H/^{12}C_2{}^{1}H$ data.

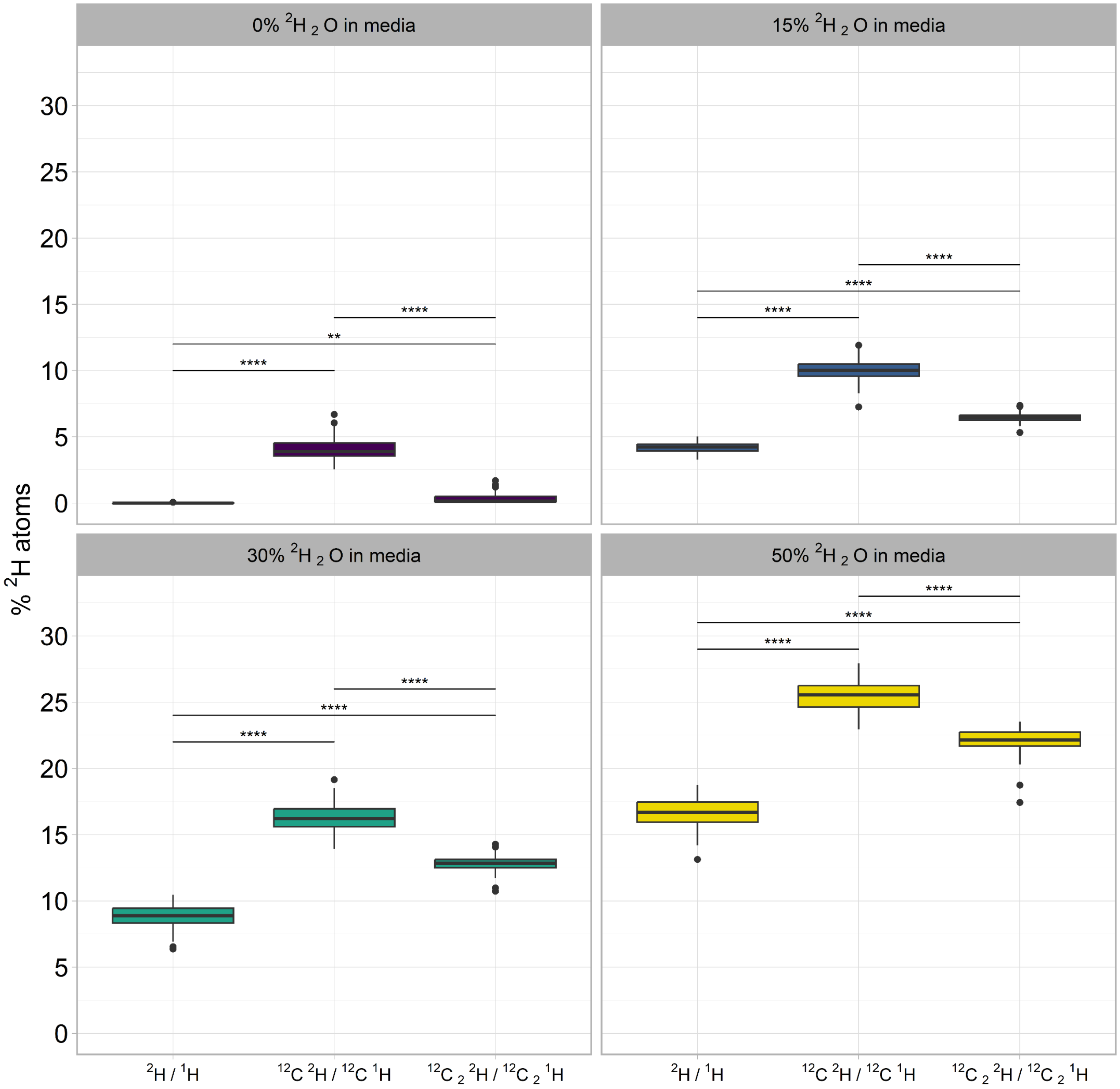


Fig. S8. Percent $^{2}$H as measured by the different atomic mass ratios using NanoSIMS across the four heavy water incubations. All statistically differences are shown: ** = p-value < 5.0 x $10^{-3}$, **** = p-value < 1.0 x$10^{-10}$.

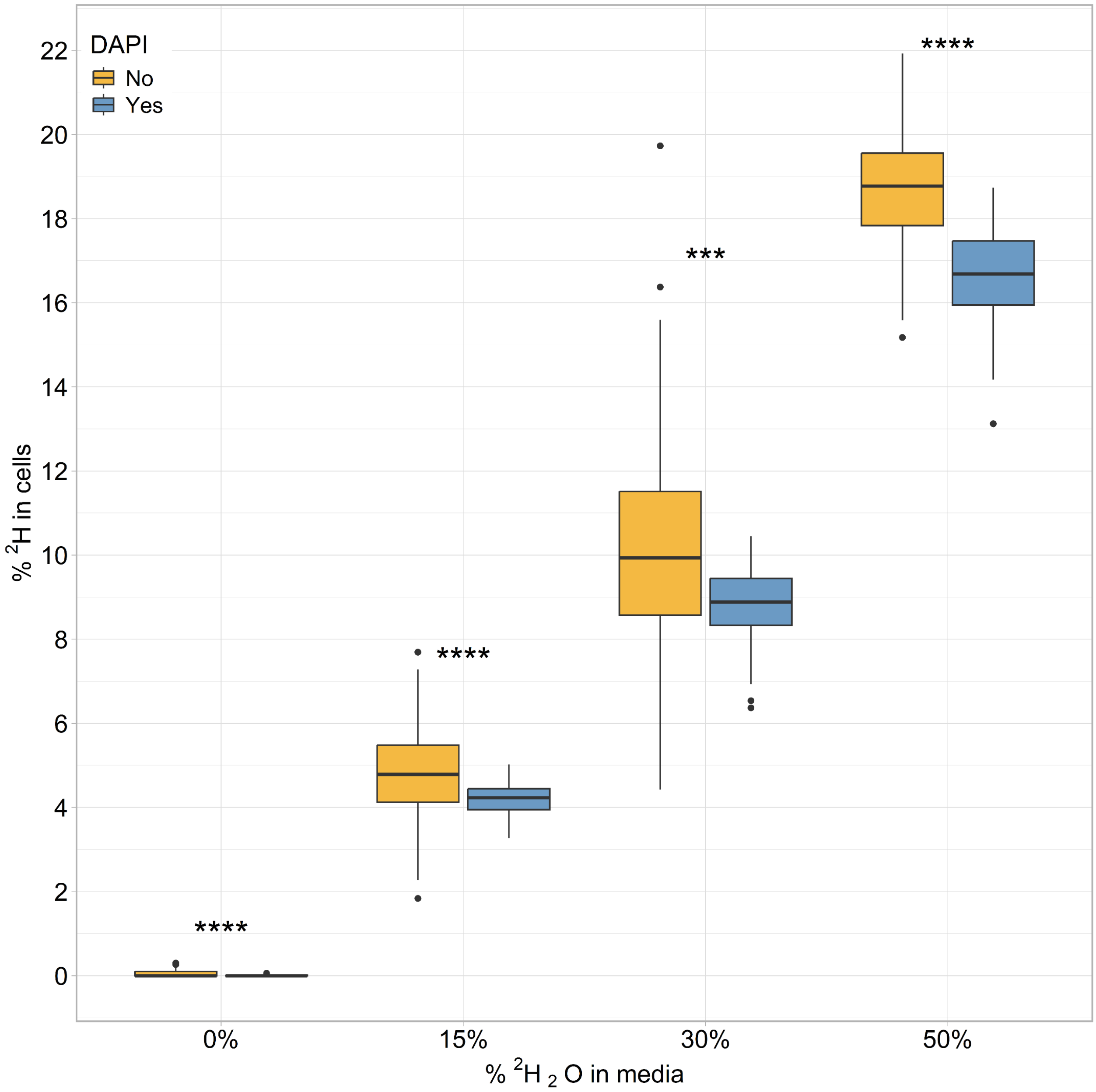


Fig. S9. NanoSIMS measurements of mass $^2H/H$ for *E. coli* cells either stained with DAPI or not. Cells that had not been stained with DAPI showed a statistically higher $^2H$ content than cells that had been treated with DAPI (*** = p-value < $1.0\times10^{-4}$, **** = p-value < $1.0\times10^{-5}$)

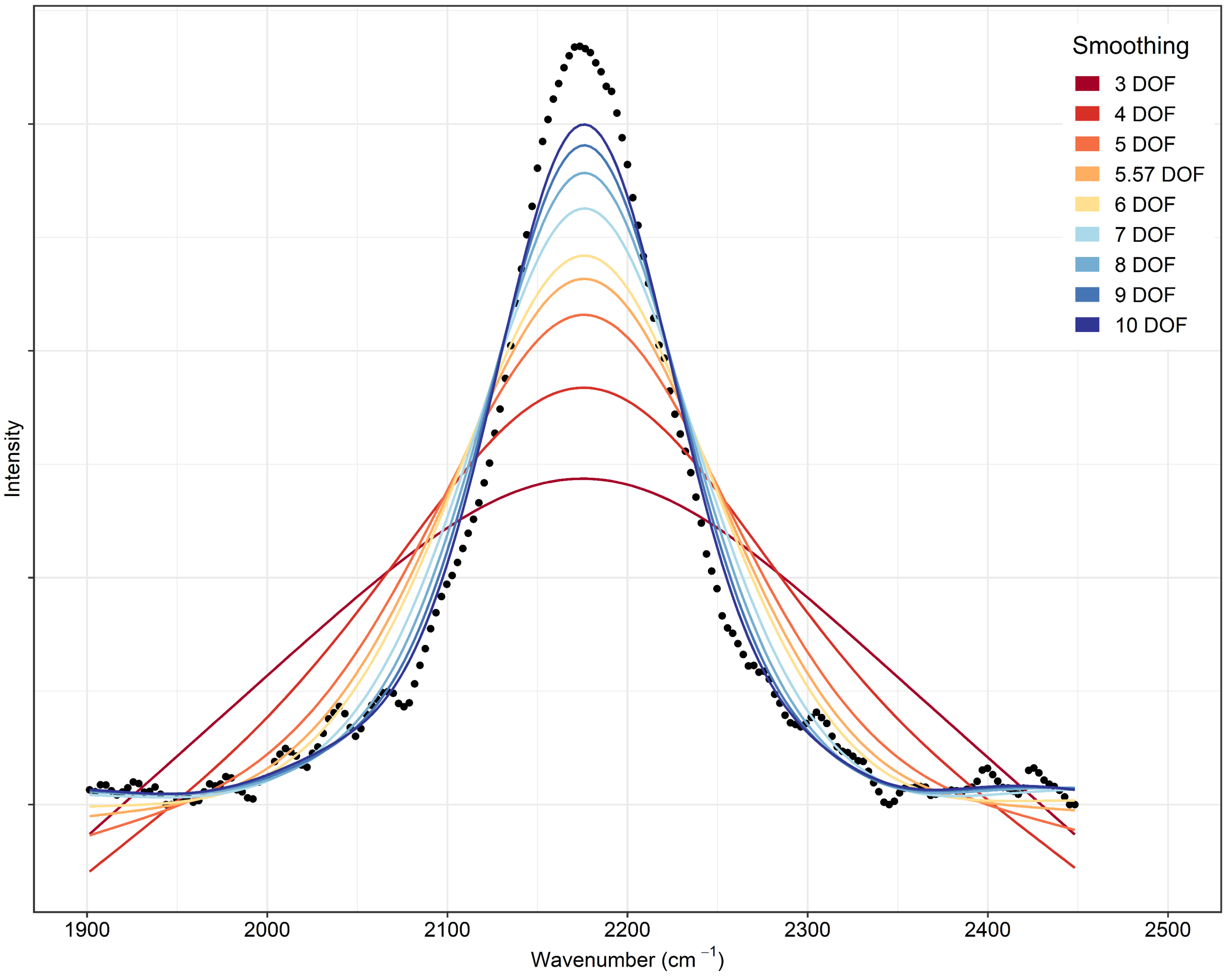


Fig. S10. Comparison of degrees of freedom (DOF) for the spline smoothing function of as applied to the $^{12}C^{2}H$ peak in a spectra obtained for a cell from the 50% $^{2}H_2O$ incubation. The higher the DOF, the better the fit to the data, but the further constrained the wavenumber range for the 2$^{nd}$ derivative becomes (see Figs S10 and S11).

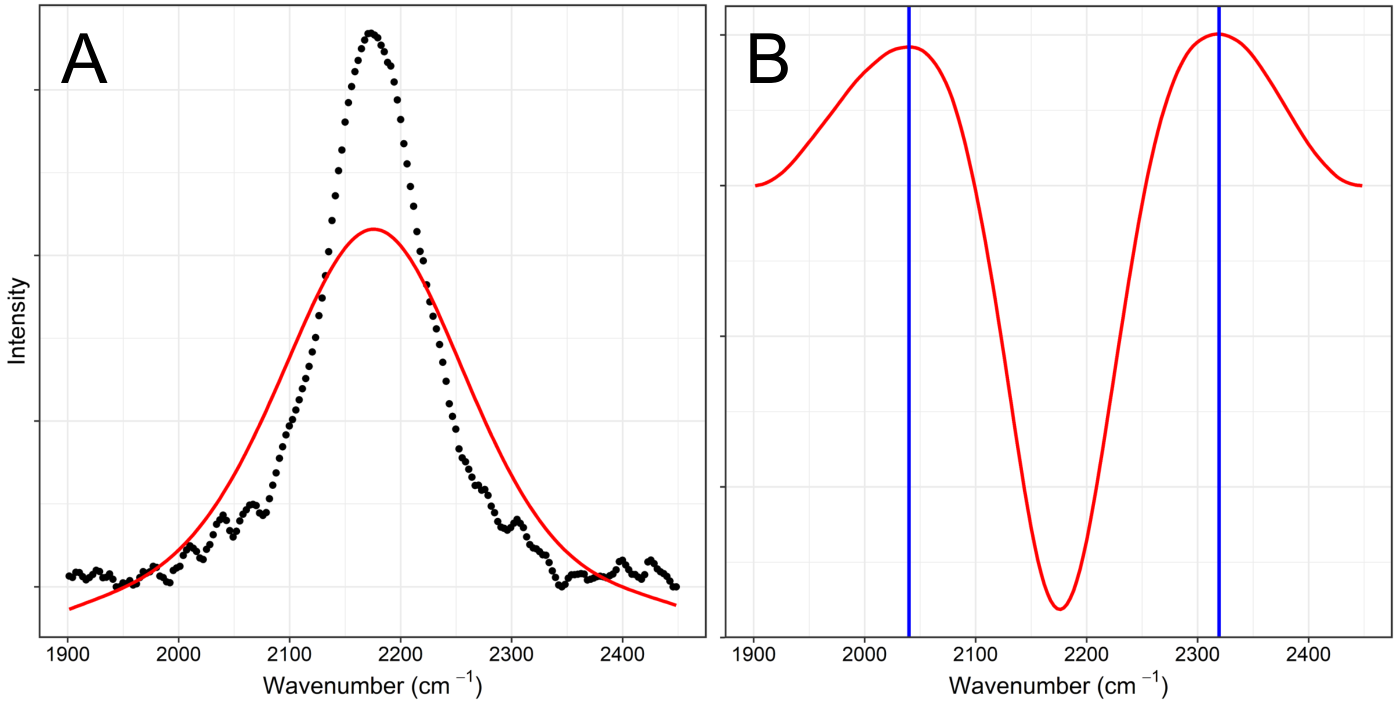


Fig. S11. Statistical evaluation of wavenumber range for 50% $^2H_2O$ incubation spectra. **A** Processed spectra (black dots) with five degrees of freedom (DOF) spline smoothing (red line). **B** Calculation of the 2nd derivative for the red line shown in panel **A**. The inflection points (blue vertical lines) are the max values of the curve on either side of the lowest value for the 2nd derivative of the curve and indicate the wavenumbers as determined by the 2nd derivative.

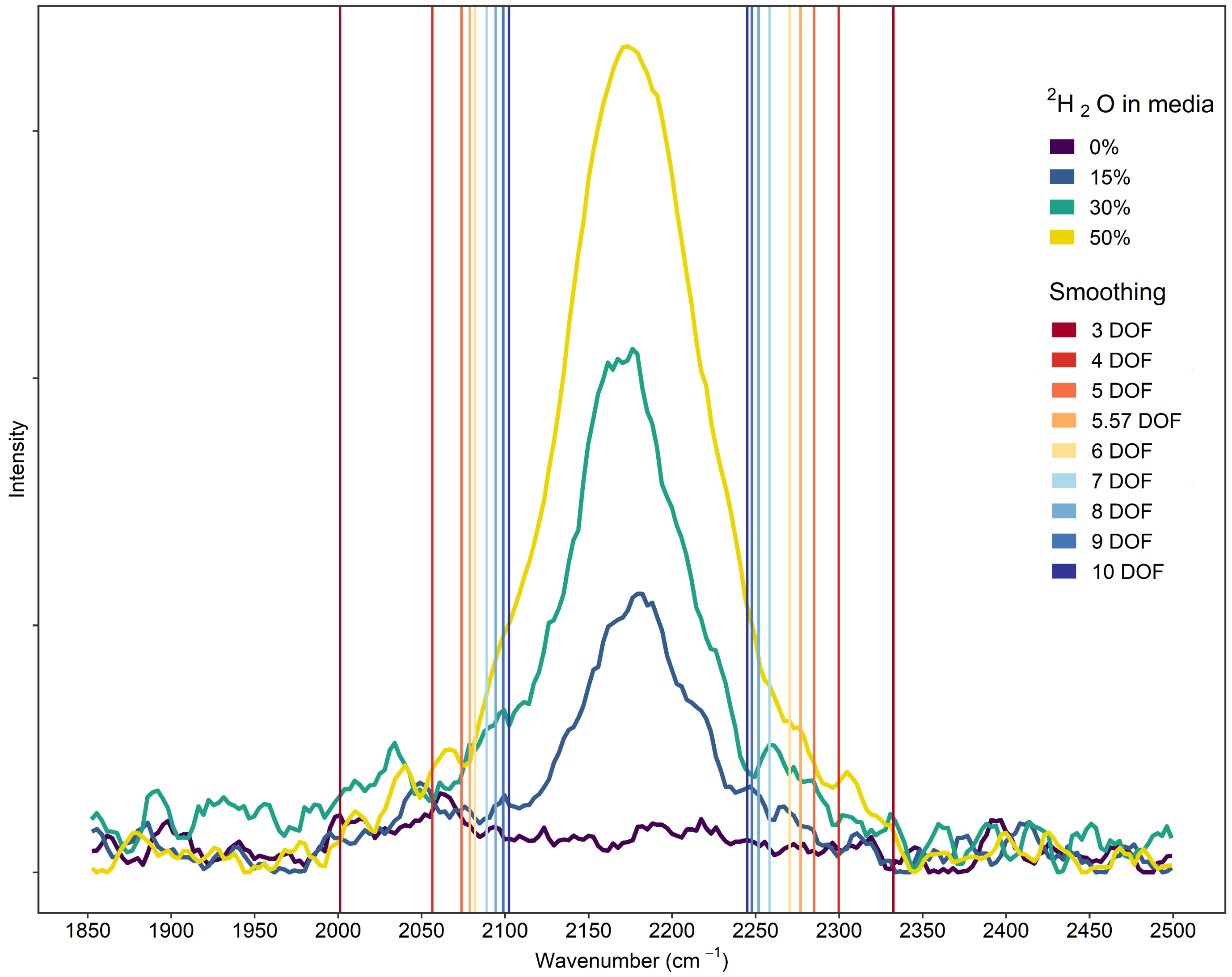


Fig. S12. Plot showing the wavenumber ranges (shown as vertical lines) as calculated by the 2nd derivative for the different degrees of freedom (DOF) smoothing. Example spectra for cells incubated in 0%, 15%, 30%, and 50% $^2H_2O$ are shown for reference.

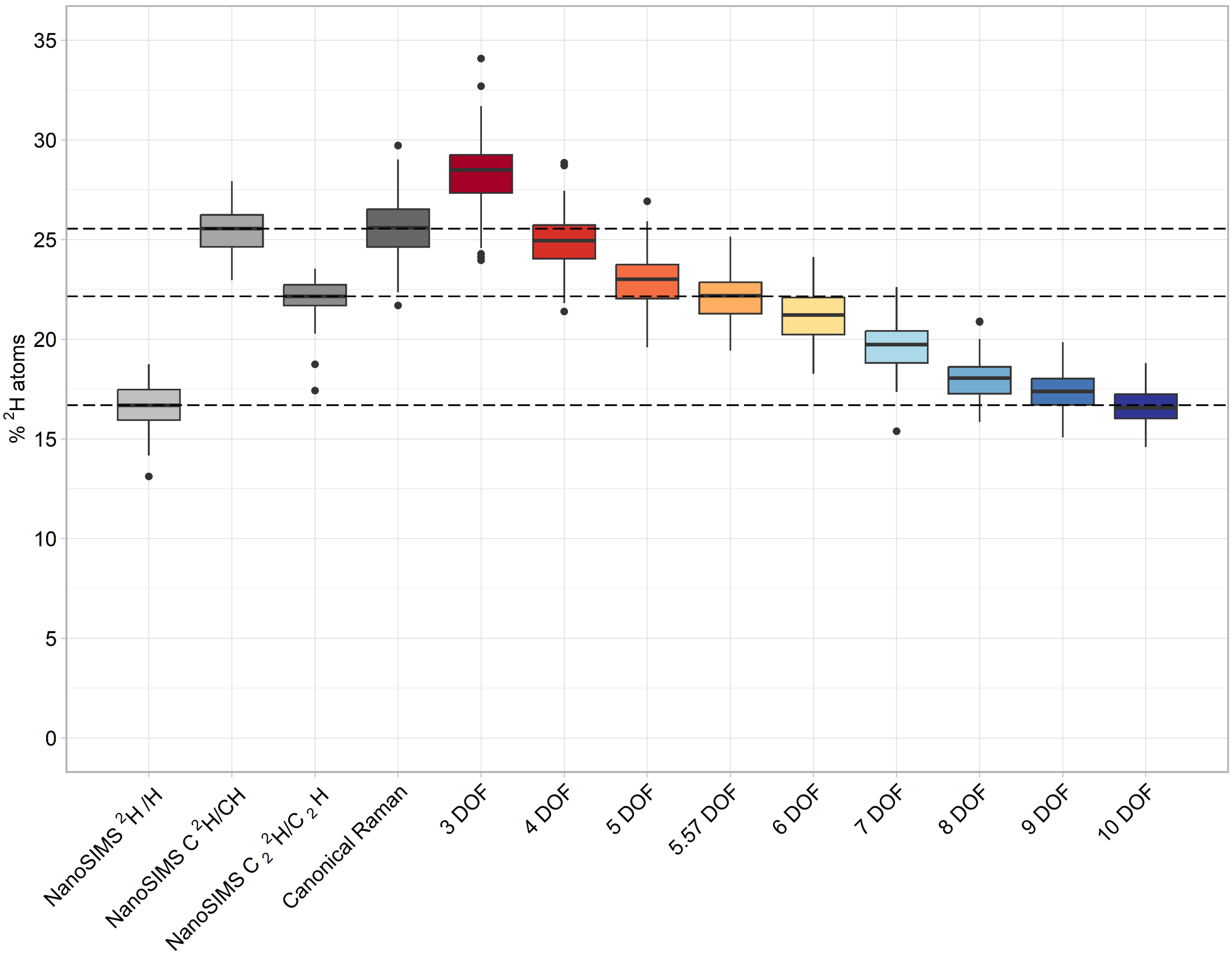


Fig. S13. Comparison of the atom percent of $^2H$ in cells from the 50% $^2H_2O$ incubation as determined by NanoSIMS and Raman analysis using the different wavenumber ranges calculated for the different degrees of freedom (DOF). Dashed horizontal lines indicate each DOF that best matches the respective NanoSIMS mass fraction.

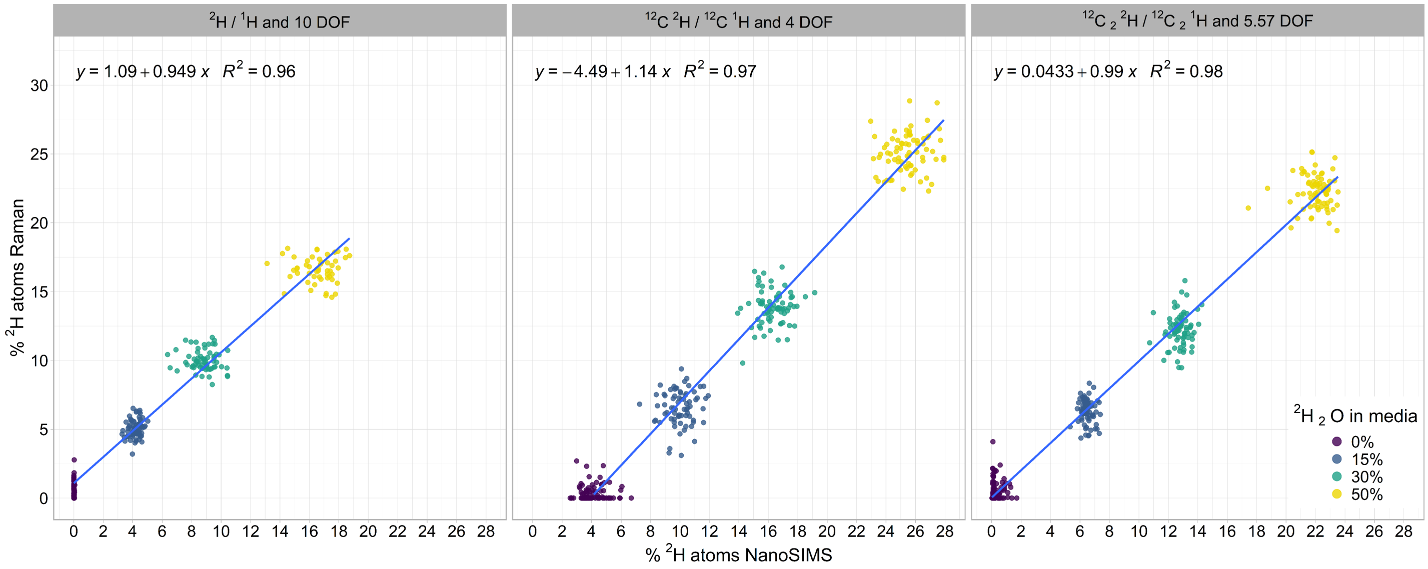


Fig. S14. Single cell comparison of the $^2$H content of cells as measured by Raman using the 2nd derivative wavenumber ranges to NanoSIMS using specific mass ratios. Each dot represents a single cell analyzed with both techniques for different $^2H_2O$ concentrations in the culture medium. The linear model equation and fit (blue line) is shown for each comparison of Raman to NanoSIMS regarding the specific mass ratios.

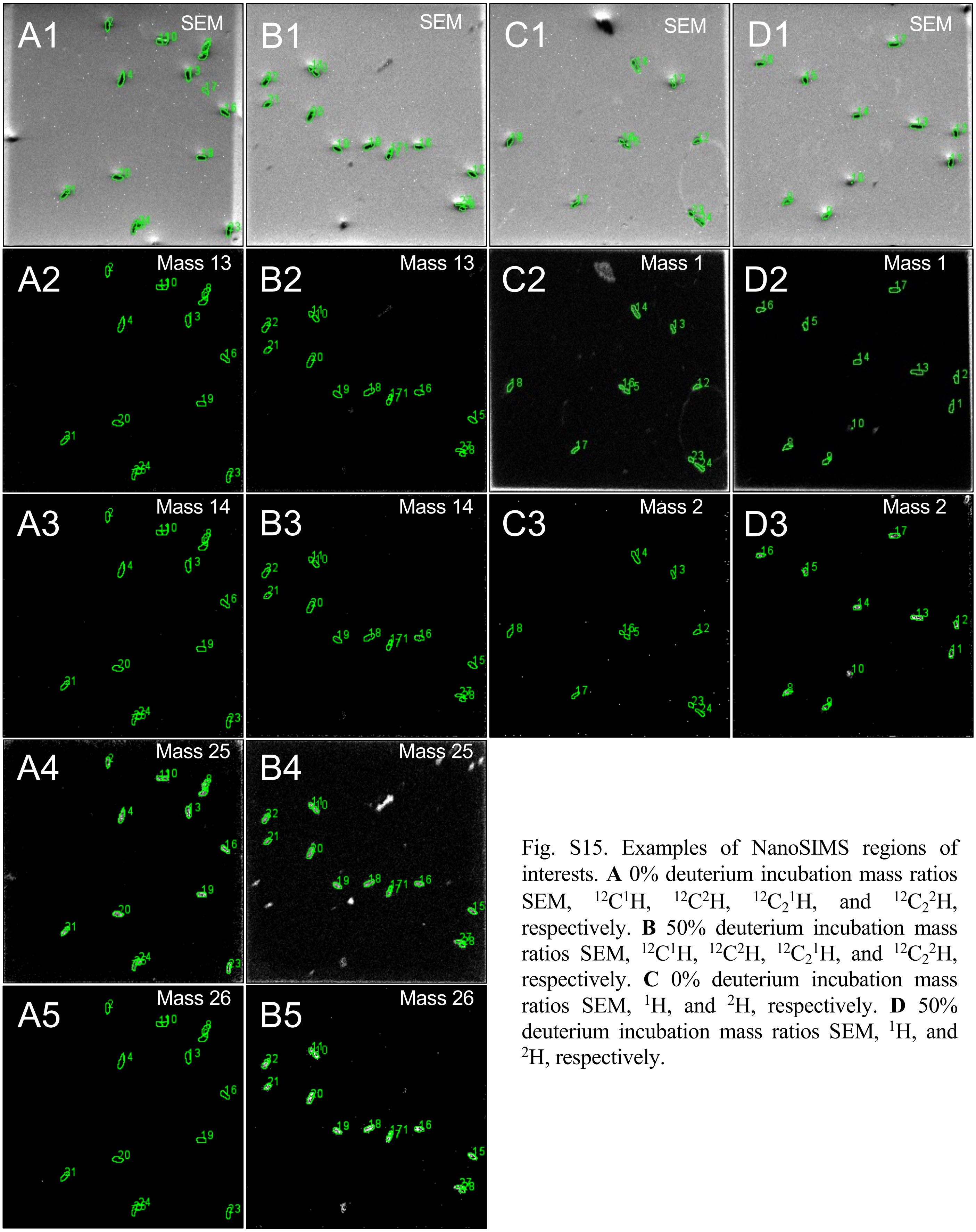


Fig. S15. Examples of NanoSIMS regions of interests. **A** 0% deuterium incubation mass ratios SEM, $^{12}C^{1}H$, $^{12}C^{2}H$, $^{12}C_2{}^{1}H$, and $^{12}C_2{}^{2}H$, respectively. **B** 50% deuterium incubation mass ratios SEM, $^{12}C^{1}H$, $^{12}C^{2}H$, $^{12}C_2{}^{1}H$, and $^{12}C_2{}^{2}H$, respectively. **C** 0% deuterium incubation mass ratios SEM, $^{1}H$, and $^{2}H$, respectively. **D** 50% deuterium incubation mass ratios SEM, $^{1}H$, and $^{2}H$, respectively.

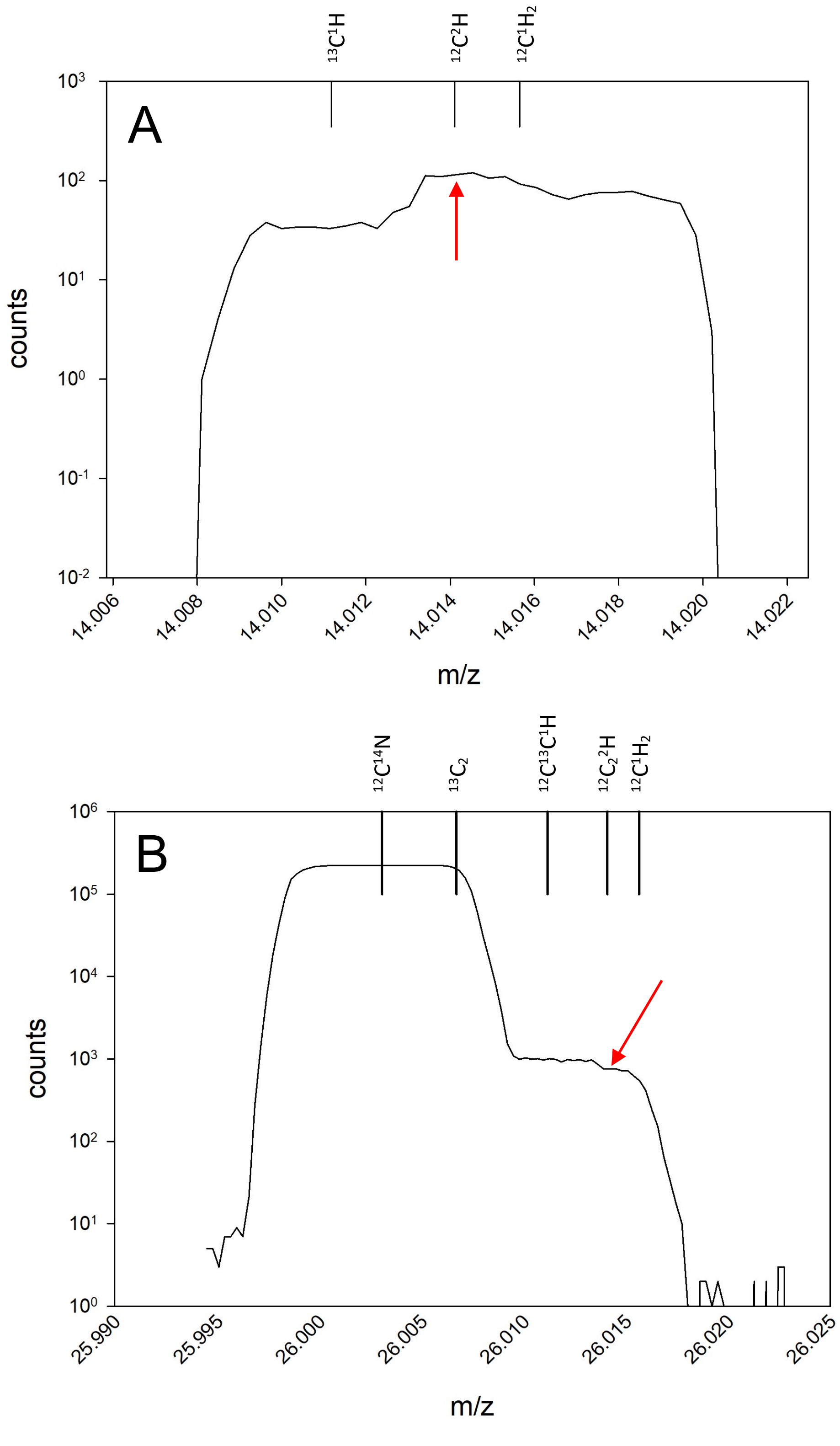


Fig. S16. Optimization of the deflector settings in NanoSIMS to minimize isobaric interferences. Mass spectra of (**A**) m/z 14 and (**B**) m/z 26 show isobaric interferences considered for deflector tuning. Red arrows indicate the peaks of interest for $^{2}H$ incorporation.
